# Photo-downregulation of SIRT4 mitigates aging in mice by enhancing H3K9ac via fatty acid metabolism

**DOI:** 10.64898/2026.04.07.717004

**Authors:** Fangqing Deng, Rong Yang, Xu Li, Jinyun Niu, Zibo Gao, Monian Wang, Yang Liu, Lihua Yang, Huifang Liu, Yingchun Yang, Zhaoxiang Yu, Lianbing Zhang

## Abstract

As organisms age, mitochondrial metabolic activity declines, and disrupted gene expression regulation mediated by histone acetylation induces the emergence of senescent physiological phenotypes in tissues. In this study, we found that periodic exposure to red light significantly increased histone H3 Lys9 acetylation (H3K9ac) levels in the tissues and organs of aged mice. Following red light exposure, silent information regulation factor 4 (SIRT4) protein levels in keratinocytes were notably reduced, whereas glycolysis, fatty acid metabolism, and the tricarboxylic acid (TCA) cycle were significantly activated in keratinocytes. The reduction in mitochondrial SIRT4 levels enhances the acetylation of mitochondrial metabolic proteins, particularly malonyl-CoA decarboxylase (MCD), a potent inhibitor of the key rate-limiting enzyme carnitine palmitoyltransferase 1A (CPT1A) in fatty acid oxidation. This process promotes mitochondrial fatty acid oxidation and TCA cycle. Additionally, the decrease in SIRT4 activates SIRT1 through feedback mechanisms, thereby alleviating its inhibition on PPAR-α in senescent keratinocytes and comprehensively activating the expression of genes related to lipid metabolism. This lipid metabolism activation ultimately facilitates the accumulation of acetyl-CoA within keratinocytes, increases H3K9ac levels, and reshapes the expression patterns of senescence-related genes. Eventually, cellular aging is effectively mitigated by the synergistic regulation of metabolism, inflammation, and gene expression.

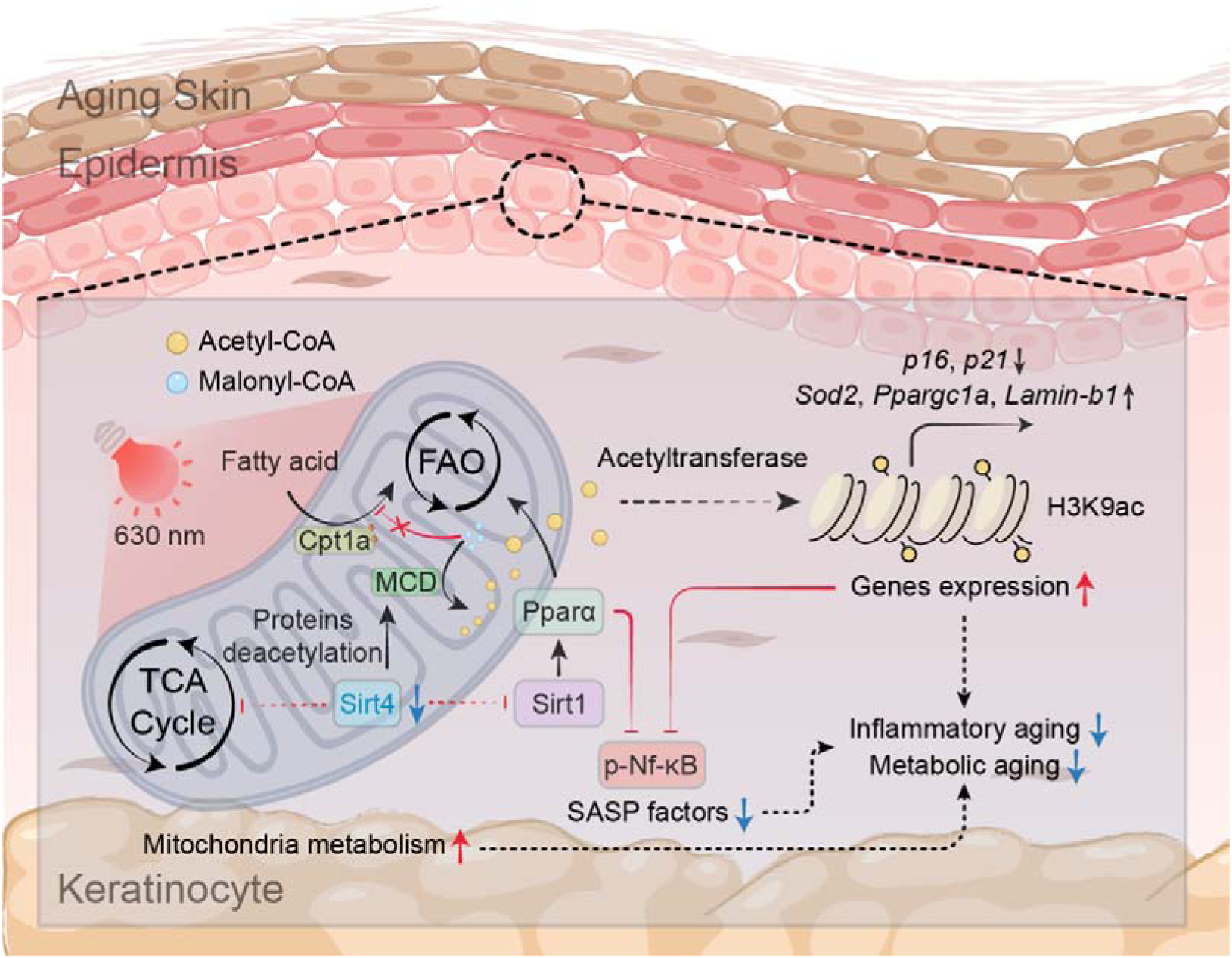

**Graphical Abstract:** **Mechanism of anti-aging action of red light:** Red light can reduce SIRT4 signalling in keratinocytes, thereby reactivating lipid metabolism and increasing levels of acetyl-CoA. This promotes histone acetylation, which in turn remodels the expression of age-related inflammatory factors and genes.

## Introduction

Aging is characterized by the onset of chronic inflammation, cardiovascular fibrosis, subcutaneous melanin accumulation, wrinkle formation, and other physiological changes^1, 2^. These alterations pose significant risks to the physical and mental wellbeing of the elderly population^3^. Therefore, it is crucial to thoroughly investigate the mechanisms of aging and develop comprehensive anti-aging strategies and interventions.

Existing research has elucidated the mechanism by which histone acetylation influences aging phenotypes through genomic expression regulation^4^. Changes in the acetylation levels at age-related histone acetylation sites directly shape cellular and tissue aging phenotypes^5, 6^. For example, elevated levels of H3K9ac, H3K56ac, and H3K27ac maintain chromatin accessibility, enhance binding of relevant transcription factors, and not only upregulate antioxidant genes (*SOD2*, *NRF2*) to suppress NF-κB-mediated inflammatory signaling and SASP aging factor secretion^7–11^, but also directly inhibit promoters of cell cycle suppressors (P16^INK4a^, P21^CIP^^1^)^12–14^. As a key metabolite and the sole substrate for histone acetylation, acetyl-CoA links cellular metabolic states to senescence phenotypes through histone acetylation^4, 15^. Under the influence of intracellular acetyltransferase and deacetylase systems, increased or decreased cellular metabolism affects histone acetylation levels by regulating acetyl-CoA content^16, 17^. This pattern of gene expression, regulated by metabolism and histone acetylation during cellular aging, provides a strategy for delaying aging phenotypes. Specifically, by increasing cellular metabolism or inhibiting histone deacetylase activity, the modification of senescence-associated histone acetylation sites can be enhanced to delay cellular senescence^18–20^.

Protein acetylation levels are regulated by deacetylases and acetyltransferases^21–23^. The activities of these enzymes are closely linked to the metabolic state of the cell^4^. In particular, with respect to protein deacetylases, three major classes of histone deacetylases (HDACs) have been identified within cells^22^. Class I and Class II HDACs are both Zn²□-dependent and primarily localized to the nucleus^24^. Class III HDACs comprise the NAD□-dependent Sirtuins (SIRT1-SIRT7) family proteins, which exhibit diverse localization patterns and possess deacetylase activity^25, 26^. Notably, Sirtuins deacetylate not only histones but also numerous non-histones, including transcription factors, metabolic enzymes, and DNA repair proteins, playing central roles in regulating metabolism, stress resistance, genomic stability, inflammation, and aging^27–29^. Among these Sirtuins, SIRT7 is expressed in the nucleolus and directly regulates ribosomal transcription, making it a key gene for cellular survival^28, 30^. While studies have clearly demonstrated that overexpression of SIRT1/6 extends lifespan in model organisms and that SIRT3/6 deficiency causes severe premature aging in mice, the anti-aging mechanisms of SIRT4/5 remain incompletely understood^30–35^. In calorie-restriction experiments, SIRT4 activity was suppressed, suggesting that SIRT4 may serve as potential feedback regulators of Sirtuins^30^. Unlike Class I and Class II HDACs, Sirtuins exhibit NAD□ dependence. Consequently, current anti-aging strategies targeting sirtuins primarily focus on elevating NAD□ levels (through supplementation of NAD□ precursors such as NMN/NR) or employing sirtuin-specific activators, which are currently popular anti-aging interventions^36^.

Distinct from the inhibitory effects of short-wavelength blue light on cellular viability and its pronounced induction of reactive oxygen species, red light possesses a robust intrinsic capacity to regulate cellular metabolism^37, 38^. As an environmental stimulus, photobiomodulation mediated by 625–635 nm red light is primarily characterized by enhanced cellular metabolism and attenuated inflammation^37, 39, 40^. In clinical settings, visible red light is routinely employed to accelerate metabolic activity in hair follicles and treat diverse cutaneous inflammatory responses^37, 41, 42^. As a noninvasive intervention, the effects of red light have also garnered public interest through devices such as ‘photorejuvenation devices’ and ‘home wearable wound light care,’ which have yielded significant social and economic benefits, underscoring their efficacy in skin repair and potential for anti-aging applications^43–46^. Notably, the metabolic remodeling capacity of red light implies a potential link to the regulation of histone acetylation. However, the relationship between red light-driven metabolic regulation and histone acetylation, as well as the mechanism by which red light regulates histone acetylation, remain unclear. We sought to understand how red light exerts these beneficial effects through the regulation of histone acetylation and its specific molecular mechanisms. Therefore, we irradiated aging mouse and cell models with red light and monitored changes in aging-related physiological indicators post-treatment to explore the potential of red light for epigenetic modification and antiaging effects.

In our study, we found that periodic exposure to red light significantly increased H3K9ac levels in the tissues and organs of aged mice. Following red light exposure, SIRT4 protein levels in keratinocytes were notably reduced, while glycolysis, fatty acid metabolism, and the tricarboxylic acid (TCA) cycle were significantly activated. Further mechanistic investigations suggested that the reduction in mitochondrial SIRT4 levels enhances the acetylation of mitochondrial metabolic proteins, particularly malonyl-CoA decarboxylase (MCD), a potent inhibitor of the key rate-limiting enzyme carnitine palmitoyltransferase 1A (CPT1A) in fatty acid oxidation. This process promotes mitochondrial fatty acid oxidation and TCA cycle activity. Additionally, the decrease in SIRT4 activates SIRT1 through feedback mechanisms, thereby alleviating its inhibition on PPAR-α in senescent keratinocytes and comprehensively activating the expression of genes related to lipid metabolism. This lipid metabolism activation ultimately facilitates the accumulation of acetyl-CoA within keratinocytes, increases H3K9ac levels, and reshapes the expression patterns of senescence-related genes. Consequently, it mitigates the dysregulation of genomic expression and excessive inflammatory activation associated with cellular senescence. Ultimately, cellular aging is effectively mitigated through the synergistic regulation of metabolism, inflammation, and genomic expression. In a word, our study reveals a novel anti-aging intervention strategy targeting SIRT4 downregulation to increase H3K9ac levels. We propose that controlling the abnormal age-related activation of SIRT4 during cellular senescence and developing SIRT4 inhibitors may enable tissue-specific senescence reversal and therapeutic interventions for metabolic diseases.

## Methods

### Cell culture and animal models

PAM212 (Cat. No. MZ-2610) cell lines were purchased from Ningbo Mingzhou Biotechnology Co., Ltd. Under non-experimental conditions, the cells were maintained in Dulbecco’s Modified Eagle’s medium (DMEM, C11995500BT, Gibco, USA). containing 10% (v/v) fetal bovine serum (FBS), 100 U/mL penicillin, and 100 mg/mL streptomycin. Prior to red light irradiation treatment, the old medium was removed, the cells were carefully rinsed with D-PBS, and the medium was replaced with phenol red-free DMEM. For both irradiation and maintenance cultures, the cells were placed in a cell culture incubator, and the temperature of the incubator was maintained at 37 °C, 80% humidity, and 5% CO2.

SPF-grade female C57BL/6 mice (4–5 weeks) were purchased from Xi’an Keao Biotechnology Co., Ltd. The mice used in this study were housed in a dedicated experimental microbarrier (HH-MMB-1, Monkey Animal Experimental Equipment Technology Co., Ltd., Suzhou, China) and provided with an unrestricted standard laboratory diet and water. The microbarrier temperature was maintained at 26 °C. When the mice reached the age of 8 months, periodic red light irradiation treatments were performed according to the light treatment protocol for mice in this study. Red-light-irradiated mice were analyzed at the end of treatment. All animal experiments involved in this study were approved by the Animal Ethics Committee of Northwestern Polytechnical University (Ethics No. 202401023).

### Model of keratinocyte senescence

2 × 10□ PAM212 cells were seeded in 6-well plates and incubated overnight in a cell culture incubator. When the cell confluency reached 50%-60%, the medium was removed, and 2 mL of PBS was added to rinse the cells. After replacing the serum-free medium with H_2_O_2_ at a concentration of 300 μM, the cells were returned to the cell culture incubator and treated for 2 h to induce premature senescence following oxidative stimulation. After H_2_O_2_ treatment, the old medium was removed, the cells were rinsed twice with 2 mL of PBS, and the complete medium was replaced and incubated for 72 h. During this period, the fluid was changed, and treatments were added according to the actual conditions. Cellular senescence was detected using SA-β-Gal staining.

### Lighting equipment and light treatment solutions

The light source used in this study was purchased from Xuzhou Aijia Electronic Science and Technology Co., Ltd., and had a rated power of 12 W. The light source was emitted by LEDs, and 96 LEDs were distributed on a planar light plate according to the standard wells of a 96-well cell culture plate. The wavelength range of the LED light source was 625 nm-635 nm. The cells and mice were irradiated in an opaque darkroom with a bottom area of 400 cm^2^, and the power density of red light received at the bottom plane of the darkroom was 0.03 J/(cm^2^·s) under rated power operating conditions. For the red light irradiation source employed in this study, the distance from the irradiation plane of the mouse (the dorsal skin of the mouse) was 10 cm, and the distance from the irradiation plane of the keratinocytes was 15 cm. In this study, four irradiation dose gradients of 0 J/cm^2^, 40 J/cm^2^, 80 J/cm^2^ and 160 J/cm^2^ were used to treat cells and mice. The irradiation treatment times for the four dose gradients were 0, 1333, 2667, and 5333 s. The irradiation treatment of the cells was carried out in a cell incubator, and the irradiation treatment experiments were conducted in a low-light, room-temperature environment. Details of the light treatments are provided in the Supplementary Material (**Fig. S1**). Unless otherwise specified, the cellular treatment dose was set at 80 J/cm^2^.

Red light irradiation of senescent mice was performed according to the following procedure: C57BL/6 mice aged 8 months were used, and all hair on the backs of the mice was removed using a depilatory cream prior to irradiation. The mice were then subjected to continuous red light treatment according to a treatment cycle of 15 days of red light treatment every two months. During each treatment cycle, the mice were irradiated with red light at a dose of 80 J/cm^2^ once daily for 15 days. Mice in the control group were subjected to the same dehairing treatment but were not red-light irradiated. Red-light-treated model mice and normal senescent mice reaching 1 and 2 years of age were euthanized with an overdose of sodium pentobarbital, and tissue samples from these senescent mice were collected for the analysis of senescence indices and related experiments. All senescent mice used in this study were female.

### RNA Sequencing and Transcriptome Analysis

2×10^6^ PAM212 in 10 cm cell culture plates with 80% cell fusion was replaced with phenol red-free DMEM medium, and then treated with red light irradiation to expose them to red light irradiation at a dose of 80 J/cm^2^. The control group underwent the same fluid change operation without any irradiation. The number of biological replicates was set to n=3. After irradiation, the cells were cultured in an incubator for 12 h. The cells were then collected using a cell scraper, centrifuged to obtain a cell precipitate, and snap frozen in liquid nitrogen. After extracting cellular RNA, the transcripts of red light-irradiated PAM212 cells were sequenced by Sangon Biotech (Shanghai) Co., Ltd. The raw image data files obtained using Illumina HiSeq™ were converted into raw sequencing sequences using CASAVA Base Calling Analysis, and the sequencing sequences were assessed for quality using FastQC. Quality-assessed transcripts were used for genome structure, expression level, differential expression, and gene enrichment analyses, and the results of the differential analysis were plotted and visualized.

### Proteomics and acetylated proteomics analysis

The same cell processing procedure used for transcriptome sequencing analysis was used to treat keratinocytes. After red light irradiation, the keratinocytes were cultured in an incubator for 24 h. Keratinocytes were collected by cell scraping, centrifuged to obtain a cell precipitate, and snap-frozen in liquid nitrogen. Total cell protein was extracted (n=3, the number of keratinocytes in each sample was more than 5×10^7^), and the total protein of keratinocytes treated with red light irradiation was obtained from Shanghai Oebiotech Co., Ltd. for proteomic analysis and acetylation modification. After the extracted proteins were enzymatically digested, LC-MS/MS was used to identify the digested peptides, and the mass spectrometry data of each sample were collected in combination with DIA for spectral matching and proteomic library construction. For acetylated proteins, acetylated peptides were first enriched by acetylation, and then the library was constructed using LC-MS/MS (4D-DIA). After library construction, the peptides were processed for the analysis of differentially expressed proteins and differentially acetylated proteins after quality control. The results of the proteomics and acetylation-modified proteomics difference visualization analyses were plotted by comparing the data of the differentially expressed proteins in the GO and KEGG databases.

### Real-time Quantitative Polymerase Chain Reaction

RNA was extracted using the FastPure Cell/Tissue Total RNA Isolation Kit (RC113-01, Vazyme Biotech, China). For PAM212 cells, after removing the culture medium, the cells were covered with lysate (500 µl buffer SRL for every 2×10^6^ cells), and the cell lysate was collected to extract RNA, according to the manufacturer’s instructions. The tissues were then frozen in liquid nitrogen and ground into a powder. A total of 20 mg of animal tissue was collected and added to 500 µL buffer SRL, and RNA was extracted according to the manufacturer’s instructions. RNA concentration was determined using a Nanodrop (NanoDrop One, Thermo Fisher, USA) after obtaining RNA. A total of 500 ng of RNA was extracted from each sample and reverse-transcribed using ABScript III RT Master Mix (RK20429, Abclonal, China) for qPCR. Finally, real-time quantitative PCR was used to detect differences in gene expression between the groups. The QPCR instrument was purchased from Bioer Technology Co., Ltd. (FQD-96A, Bioer Technology, China). Primers used for qPCR were listed in Supplementary Table S1.

### Western blot

Treated PAM212 cells or liquid nitrogen snap-frozen and ground animal tissue samples were added to an appropriate amount of RIPA lysate (ZG394617, Thermo Fisher, USA, 100 μl RIPA lysis buffer per 1 × 10□ cells or 600 μl RIPA lysis buffer per 20 mg animal tissue) to extract the total protein. Lysis was performed on ice for 30 min, during which the cells were vortexed and shaken for 10 s every 10 min. The samples were centrifuged for 15 min at 4°C and 12,000 × g. The supernatant (total protein) was aspirated into a new EP tube. The total protein concentration of the protein samples was determined using the BCA assay (P1511, Beijing Applygen Gene Technology, China). All samples were adjusted to the same concentration using RIPA lysate, and 5× SDS-PAGE buffer (RM00001, Abclonal, China) was added to each sample. The proteins were denatured by boiling in a water bath for 10 min. Sampling was performed using a gradient precast gel (ET12420Gal, ACE Biotechnology, China) from ACE Biotechnology Co., Ltd., and the sample volume was 10-20 μg of protein. Electrophoresis (constant voltage of 120 V for approximately 60–90 min) was performed using an electrophoresis chamber (PowerPac Basic, Bio-Rad, USA).

Membrane transfer was performed using a rapid membrane transfer apparatus (PB0010, Thermo Fisher, USA). Polyvinylidene fluoride (PVDF, #1620177, Bio-Rad, USA) membranes were purchased from Bio-Rad Laboratories Inc. After transfer, the PVDF membranes were blocked using a rapid blocking solution from New Cell & Molecular Biotech Co.,Ltd. The PVDF membranes were then incubated with the following primary antibodies: NF-κb (A22331, Abclonal, China), p-NF-κB (AF5875, Beyotime, China), IκB-α (A19714, Abclonal, China), Actin (AC026, Abclonal, China), LMNB1 (A1910, Abclonal, China), P16 (A24653, Abclonal, China), P21 (A27846, Abclonal, China), p-mTOR (AP0094, Abclonal, China), Glut1 (A6982, Abclonal, China), PI3K (A22467, Abclonal, China), p-AKT1 (AA329, Beyotime, China), PPAR-α (A22887 and A25296, Abclonal, China), PPAR-γ (GB11164, Servicebio, China), H3K9ac (A21107, Abclonal, China), H3K4ac (A24340, Abclonal, China), H3K27ac (F0271, Selleck, USA), H3-K9/K14/K18/K23/K27ac (A17917, Abclonal, China), H3 (A2348, Abclonal, China), mTORC1 (A8992, Abclonal, China), mTORC2 (27248-1-AP, Proteintech, China), p-S6 (AP1471, Abclonal, China), SIRT1 (A17307, Abclonal, China), SIRT4 (A7585 and A15800, Abclonal, China), and Tubulin (A12289 Abclonal, China). After incubating the membrane with the primary antibody solution at 4 °C overnight (12-16 h), the membrane was washed six times with TBST for 5 min each. The membrane was then incubated with an HRP-labelled secondary antibody of the corresponding species at room temperature for 1 h. Finally, the membrane washing operation was repeated, an appropriate amount of ECL luminescent solution was added to the PVDF membrane, and the signals of the target bands were captured using a gel-imaging system.

Western blot band intensities were quantified using Image J software. Briefly, the captured WB images were converted to grayscale, and the integrated density of target protein bands in each lane was measured using regions of interest of identical size, followed by subtraction of the background signal from adjacent areas. The corrected band intensities of the target proteins were normalized to the corresponding Actin signal, whereas H3K9ac was normalized to total H3. The mean relative expression levels of biological replicates at each time point were then calculated. For heatmap visualization, the normalized expression values of each protein across different time points were further standardized to the maximum value of that protein. Heatmaps were generated using GraphPad Prism, with color intensity representing changes in relative protein expression levels.

### Mitochondrial membrane potential assay (JC-1)

3 × 10□ PAM212 cells were seeded in 35 mm confocal specialized culture dishes, and overnight adherence was continued until cell fusion reached 50-60%. After red light treatment of PAM212 cells, changes in the mitochondrial membrane potential in keratinocytes were detected using the Enhanced Mitochondrial Membrane Potential Assay Kit (C2003S, Beyotime Biotechnology, China). Ltd. Membrane potential collapse was induced by adding CCCP (10 μM) for 20 min to the positive-control group. The JC-1 probe working solution (5 μM) was prepared in a serum-free medium and added to the experimental and control groups. The medium was removed from the

Petri dishes, and 1 mL of preformulated JC-1 working solution was added and incubated for 20 min at 37°C in the dark. The cells were gently washed with pre-warmed JC-1 Wash Buffer (provided in the kit) 3 times for 5 min each. Phenol Red-free medium (1 mL) containing 10% FBS was added, and fluorescence imaging was performed using a confocal microscope (SP8, Leica, Germany). Microplate reader detection was performed in a 96-well plate, and staining was performed as described above. The optical density (OD) values of the different treatment groups at 488 nm (monomers, green) and 561 nm (aggregates, red) were determined.

### Cell activity, migration and proliferation assays

Changes in cell viability were detected using the CCK-8 assay (C6005, NCM Biotech, China) after treating the cells with red light irradiation. 5×10^3^ PAM212 cells were seeded in 96-well plates, and the medium was changed to phenol red-free complete DMEM after overnight wall attachment. The keratinocytes were treated with red light irradiation, and 10 μl of CCK-8 solution was added to each well after red light treatment. The OD value of the cell cultures at 450 nm was measured using a microplate reader after incubation at 37 □ for 1 h.

Changes in the migration ability of the cells were detected using a Wound Healing Assay. PAM212 cells (10^6^) were seeded in 6-well plates and incubated at 37 °C until a 90%-100% cell fusion rate was achieved. The cell layer was scraped vertically using a sterile 200 μl pipette tip to form a straight line of scratches. The cells were then gently rinsed three times with PBS to remove the shed cells, and the medium was replaced with phenol red-free DMEM containing 1% FBS. The initial state of the scratch at 0 h was recorded using an optical microscope. The cells were treated with red light irradiation, and the scratch was photographed again at the same location after 24 h of cell migration to observe the filling of the scratch by cell migration.

Changes in cell proliferative capacity were detected by cell counting. After inoculation of 2×10^5^ PAM212 cells in 6-well plates and overnight culture, the phenol red-free medium was changed, and the cells were irradiated with red light. Adherent PAM212 cells were digested with trypsin and collected at 0h, 6h, 12h, 18h, and 24h after the end of red light treatment. The number of PAM212 cells in the control and red light-treated groups at these time points was counted using a cell counter (Countstar Mira BF, Shanghai Ruiyu Biotech Co., Ltd., China).

### Chromatin immunoprecipitation–qPCR

Chromatin immunoprecipitation (ChIP) was performed using the Chromatin Immunoprecipitation Kit (RK20258, Abclone, China) according to the manufacturer’s instructions. Briefly, chromatin was prepared from treated cells and immunoprecipitated using an anti-H3K9ac antibody H3K9ac (A21107, Abclonal, China). A fraction of sheared chromatin from each sample was retained as input DNA to normalise ChIP enrichment, and a parallel immunoprecipitation using normal IgG was included as a negative control to estimate nonspecific background binding. Purified DNA from the input, H3K9ac-immunoprecipitated, and IgG control samples was obtained using a DNA purification kit (RK30100, Abclone, China) and subjected to quantitative PCR analysis. The enrichment of H3K9ac at target promoter regions was calculated after normalisation to the corresponding input DNA and correction against the IgG control. The RPL30 locus was amplified in parallel as an internal control locus to monitor ChIP-qPCR performance and inter-sample consistency. ChIP-qPCR primers are listed in **Supplementary Table S2**.

### Metabolite detection

After the cells were treated with red light irradiation, the intracellular levels of glucose (S0202S), triglycerides (S0219S), fatty acids (S02019S) were detected using metabolite assay kits produced by Beyotime Biotechnology Co., Ltd, lactate (D799851) and pyruvate (D799450) were detected using metabolite assay kits produced by Sangon Biotechnology Co., Ltd. 2×10^6^ PAM212 cells were seeded in a 10 cm cell culture dish and cultured until cell confluency reached 80%. The medium was replaced with phenol red-free DMEM, and the cells were irradiated with graded doses of red light. After irradiation, the cells were digested with trypsin and counted at 0h and 12h after irradiation. After counting, the cell precipitates were collected, and the levels of the corresponding metabolites in the cells at 0 and 12 h after exposure to different doses of red light were detected according to the metabolite extraction and assay procedure of the corresponding metabolite assay kits.

### NADPH, NADH and GSH content test

The 2 × 10^6^ PAM cells precipitates were collected according to the cell treatment and sample collection procedure in the metabolite assay experiment, and the content of NADPH, NAD^+^, and NADPH+NAD^+^ in the cells after red light treatment was measured using an NADPH assay kit (S0179, Beyotime Biotechnology, China). Similarly, NADH, NAD□, and NADH + NAD□ levels in the red light-treated cells were measured using an NADH assay kit (S0176S, Beyotime Biotechnology, China). The GSH and GSSG contents in the cells after red light treatment were measured using a GSH content assay kit (S0053, Beyotime Biotechnology, China).

### Detection of ATP content

The effects of different doses of red-light irradiation on cellular ATP production after cell treatment were determined using an ATP content assay kit (S0027, Beyotime Biotechnology, China). Ltd. Immediately after red light irradiation, cell lysate (200 µl per well of a 6-well plate provided in the kit) was added to the Petri dish and lysed on ice for 5 min. The cell lysate was collected, centrifuged at 12000 × g for 5 min at 4 °C, and the cell supernatant was collected. Next, 20 µL of the cell supernatant to be tested and 100 µL of ATP assay working solution were added to an opaque black 96-well plate, and 560 nm emitted light was detected using a multimode microplate reader (SpectraMax M5, Molecular Devices, USA). The total protein concentration in the sample was determined simultaneously using the BCA method: ATP content (nmol/mg protein) = sample ATP concentration/protein concentration.

### **β**-Galactosidase staining

Senescent cells were stained for galactosidase using a β-galactosidase Staining Kit (G1125-100ML, Servicebio, China) from Wuhan Servicebio Technology Co., Ltd. The cell culture medium was removed from the treated cells in 6-well plates, the cells were washed twice with PBS, 1 mL of β-galactosidase staining fixative was added, and the cells were fixed at room temperature for 15 min. The cells were rinsed three times with PBS for 2 min each, incubated with β-galactosidase staining working solution, and incubated overnight at 37 °C. The cells were observed under a microscope, and images of the galactosidase staining were captured.

### Cell immunofluorescence assay

2 × 10□ PAM212 cells were seeded onto confocal focusing dishes, allowed to adhere overnight, and cultured until 50%-60% cell fusion. The cells were treated with red light and incubated for 12 h. The medium was removed, and the cells were washed twice with PBS. Cells were then fixed with 4% paraformaldehyde for 15 min at room temperature and washed with PBS three times for 5 min each. PAM212 cells were permeabilized for 10 min at room temperature using an immunostaining permeabilizing solution (P0095, Beyotime Biotechnology Co. Ltd, China), and the PBS washing step was repeated. The blocking solution was added and incubated at 37 °C for 2 h, after which it was removed and incubated with a primary antibody. The primary antibodies (SIRT4, PPAR-α, p16, and H3K9ac) were diluted 1:100-1:200 and incubated overnight at 4 °C according to the actual conditions. The washing step was repeated, and the cells were incubated with a fluorescent secondary antibody (FITC/Cy3, 1:200) produced by Wuhan Abclonal Biotech Co., Ltd for 1h in the dark at 37 °C under a seal. After removing the secondary antibody, the cells were stained with 10 µM DAPI (BL105A, Biosharp, China) for 5 min and washed again after removing the DAPI staining solution. Finally, a fluorescent anti-quencher (CR2411131, Servicebio, China) was added, and the fluorescence signal of the cells was detected using confocal microscopy.

### Tissue immunofluorescence staining and immunohistochemical staining

Mouse tissue samples were fixed in 4% paraformaldehyde (PFA) at room temperature and embedded in paraffin. Sections of 5 µm thickness were obtained by paraffin sectioning of mouse tissues using a paraffin slicer. After dewaxing and hydration, mouse tissues were stained for PPAR-α, PPAR-γ, H3K9ac, p-NF-κB, SIRT1, SIRT4, P16, and other proteins according to the incubation procedures described above for the primary and fluorescent secondary antibodies in the cellular immunofluorescence assay. PI3K and Glut1 proteins were incubated with HRP-labelled secondary antibodies for immunohistochemical staining. After completion of tissue immunofluorescence and immunohistochemical staining, the slices were sealed with neutral resin and imaged using a tissue scanner (Science, Winmedic Tech. Co., Ltd., China) and a fluorescent tissue scanner (Pannoramic MIDI, 3DHISTECH, Hungary). Immunofluorescence images of tissue sections were quantified using ImageJ software. Images were imported into ImageJ, and regions of interest (ROIs) with the same size or corresponding anatomical areas were selected based on DAPI staining or tissue morphology. After background subtraction, the target fluorescence channel was analyzed using a consistent threshold across all groups to identify positive signals. The fluorescence signal was quantified by measuring the mean fluorescence intensity, integrated density, or the percentage of positive fluorescence area within each ROI. Multiple randomly selected non-overlapping fields were analyzed for each animal, and the average value was used as the final quantitative result for that sample.

### Bioinformatics Analysis

The visualization results of the bioinformatics analysis presented in this study were obtained using the bioinformatics cloud platform of Shanghai Oebiotech Co., Ltd. The raw transcriptomic, proteomic, and acetylation-modified proteomic data of red light-treated keratinocytes were imported into the KEGG analysis, Volcano Diagram, and Chord Diagram templates of differentially expressed genes and differentially expressed proteins of the Oebiotech Cloud Platform, according to the format required by the platform, to obtain the visualized analysis results.

For transcriptomic, proteomic, and acetyl-proteomic datasets, multiple-testing correction was performed using the Benjamini–Hochberg false-discovery-rate procedure. Differentially expressed genes or proteins were identified based on adjusted *P* values or *Q* values together with fold-change thresholds. Pathway-enrichment analyses were also evaluated using adjusted *P* values or *Q* values, and pathways with *Q* value < 0.05 were considered significantly enriched.

### Gene silencing of SIRT4

10^5^ PAM212 cells were seeded into 12-well plates (medium volume of 900 µL per well) and allowed to attach to the wall overnight. When cell fusion reached 30%-50%, the fresh medium containing the transfection reagent was replaced (810 µL complete medium + 90 µL transfection reagent). The transfection reagent was divided into solutions A and B. Solution A contained 25 pmol si-SIRT4 (50 µM, 5 µL), 50 µL Opti-MEM™ I Reduced Serum Medium (31985062, Thermo Fisher, USA), and solution B contained 50 µL Opti-MEM, 3 µL Lipofectamine RNAiMAX Transfection Reagent (13778075, Thermo Fisher, USA). The transfection complex was prepared by adding solution A to solution B, mixing well, and incubating for 5-10 min at room temperature. PAM212 cells cultured with the transfection complex were incubated at 37 °C in a 5% CO2 atmosphere for 96 h. Samples were collected at 48 h to detect the level of SIRT4 gene silencing (Qpcr) and at 96 h to detect the level of SIRT4 protein (Western blot, cellular immunofluorescence).

### PET/CT in vivo metabolic assay

Metabolic levels were measured using PET/CT in mice (2Y+R) 2 years after periodic red light treatment. A metabolically active positive control group of 4-month-old (4M) mice was used. The control group consisted of 2-year-old mice (2Y) without periodic red light treatment. After overnight fasting, the mice in the 4M, 2Y, and 2Y+R groups were weighed and injected with the corresponding dose of the radiotracer ¹□F-FDG at a dose of 4 MBq/kg. After the tracer entered the blood circulation, the mice were secured on the contrast bed with adhesive tape and visualized using a PET-CT meter to develop images. The effects of cyclic red light irradiation on the metabolism of mice were determined by analyzing the amount of radioactivity in the 4M, 2Y, and 2Y+R groups using PET-CT.

### Cellular reactive oxygen species assay (DCFH/Mito-sox)

For intracellular ROS detection, we used DCFH-DA and Mito-SOX probes to detect cytoplasmic reactive oxygen species and mitochondrial reactive oxygen species in cells, respectively. For cytoplasmic ROS detection, PAM212 cells were incubated with 10 μM DCFH-DA probe for 0.5 h at 37 °C after red light irradiation. Fluorescence imaging was performed using a fluorescence microscope (CKX53SF, Olympus, Japan). The fluorescence intensity of the DCFH-DA probe in the cells was measured using a multimode microplate reader (SpectraMax M5, Molecular Devices, USA). For mitochondrial ROS detection, cells were incubated with 5 μM Mito-SOX for 15 min, and the fluorescence intensity of the Mito-SOX fluorescent probe in the cells was characterized using a fluorescence confocal microscope and a Multi-Mode Microplate Reader.

### DHE tissue reactive oxygen species assay

After snap-freezing the dorsal skin tissues of the mice in liquid nitrogen, Service Biotechnology Co., Ltd. prepared frozen sections of the skin tissues, performed ROS fluorescence staining (using Dihydroethidium, DHE), and conducted imaging analysis using a fluorescent tissue scanner.

### H&E staining

Normal and scarred skin of mice were treated with 4% PFA and embedded in paraffin. The treated skin sample slices (5 µm) were subjected to H&E staining after deparaffinization and rehydration. Imaging was performed using a tissue slice scanner (Science, Winmedic Tech, China).

### Statistics and reproducibility

The blots presented in this study were performed in at least three replicates, and a minimum of three fields of fluorescence images were collected for fluorescence experiments. Statistical analyses were performed using GraphPad Prism. Data are presented as mean ± SEM unless otherwise stated. The number of independent biological replicates is indicated in the corresponding figure legends. For comparisons between two groups, an unpaired two-tailed Student’s *t*-test was used when the data met the assumptions of normality and equal variance. For comparisons among multiple groups, one-way ANOVA was used followed by Tukey’s multiple-comparison test for comparisons among all groups or Dunnett’s multiple-comparison test when multiple experimental groups were compared with a single control group. Normality and homogeneity of variance were assessed before applying parametric tests. Statistical significance is indicated in the figures using asterisks. The significance thresholds were defined as follows: \**P* < 0.05, \*\**P* < 0.01, \*\*\**P* < 0.001, and \*\*\*\**P* < 0.0001.

## Result

### 1. Periodic red light irradiation recovers histone acetylation and metabolism in the skin of naturally aging mice

The aging phenotype of the skin is jointly regulated by genetic and epigenetic factors^47^. However, the relationship between red light-driven metabolic regulation, histone acetylation, and skin aging has not yet been elucidated. We observed that red light (625-635 nm) irradiation at doses ranging from 0 to 160 J/cm² enhanced keratinocyte activity, with a 20% increase in cellular activity at 80 J/cm² **(Fig 1a** and **Fig S1a)**. Simultaneously, within the 0–160 J/cm² range, the acetylation levels of histone H3, particularly H3K9ac, exhibited a progressive increase as the red light irradiation dose escalated **(Fig 1b** and **S1b)**. Following treatment with 80 J/cm² red light irradiation, we observed reduced β-galactosidase activity in H□O□-induced oxidative senescence keratinocytes **(Fig 1c)**, alongside decreased expression of the cell cycle arrest proteins P16 and P21. In contrast, nuclear Lamin B1 (LAMB1), which is associated with cell proliferation, showed increased expression (**Fig S1c**). The enrichment levels of H3K9ac in the promoter regions of the anti-aging related genes *SOD2*, peroxisome proliferator-activated receptor gamma coactivator 1-alpha (*Ppargc1a*), and *LAMB1* were significantly increased (**Fig 1d-h** and **Fig S1d**). Furthermore, the expression profile of SASP inflammatory factors associated with aging is remodelled, with downregulation of cytokines such as IL-6 and IL-1α **(Fig S1e)**. These findings preliminarily demonstrate that a specific dose of red light promotes histone acetylation at the H3K9 site. As a key histone acetylation site regulating cellular inflammation and senescence, this may explain how red light alleviates oxidative senescence in keratinocytes.

**Figure 1.**
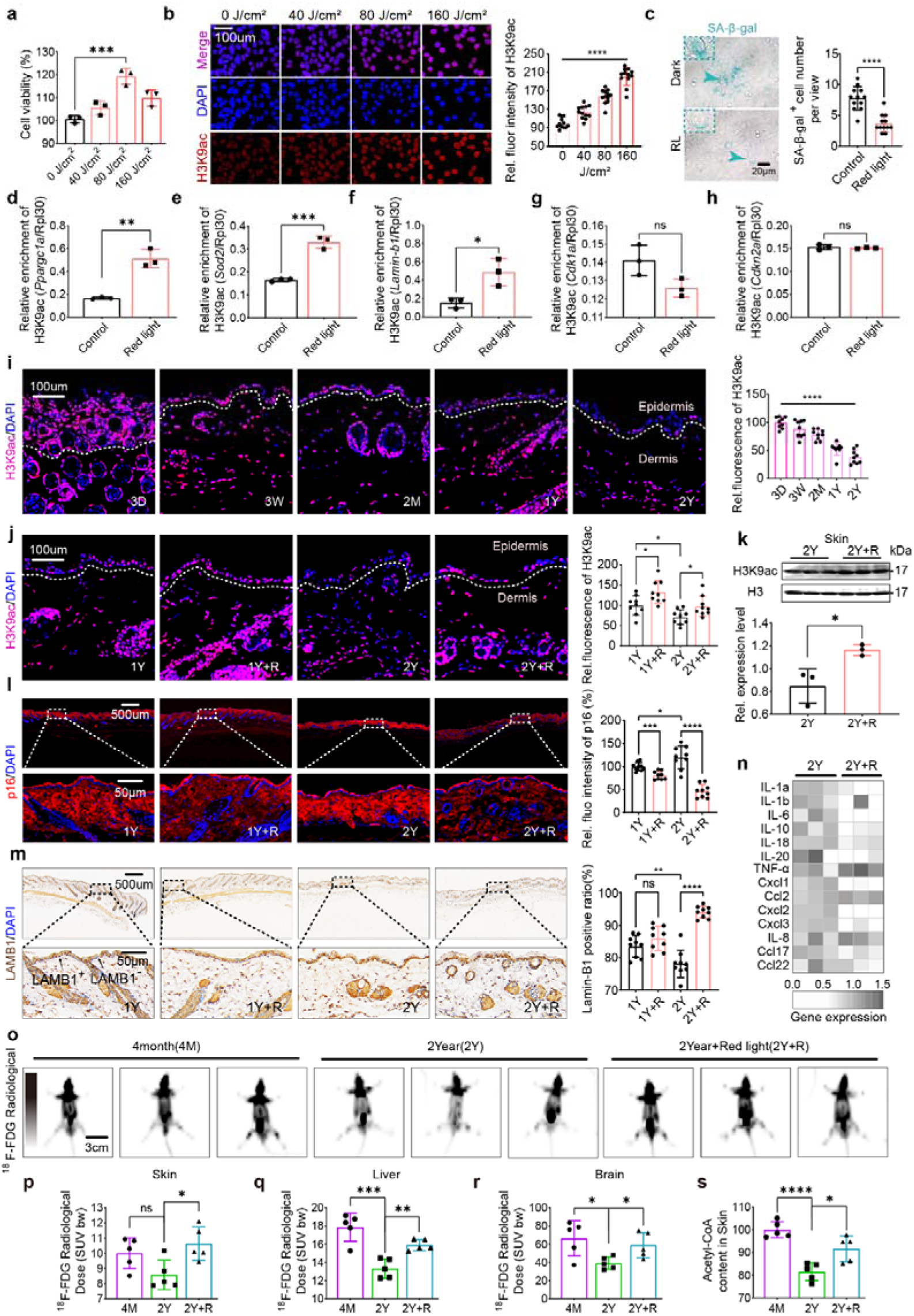
Periodic red light irradiation treatment leads to increased H3K9ac and metabolism in skin tissues of a senescent mouse model. **a**, CCK-8 analysis of keratinocyte activity after red light treatment, n=3. b, Fluorescence images of H3K9ac protein in keratinocytes after treatment with different doses of red light, ANOVA was used to analyse the significant differences in b. c, Image of β-galactosidase staining in senescent keratinocyte model after red light irradiation treatment. d-h, ChIP-qPCR experiments were conducted to assess the enrichment of H3K9ac in the promoter regions of *Ppargc1a* (d), *Sod2* (e), *Lamin-b1* (f), *Cdkn1a* (g) and *Cdkn2a* (h). Following ChIP experiments using the H3K9ac antibody or control IgG, qPCR analysis was performed using primers targeting the promoters of these genes, RPL30 is used as an internal standard for the calibration of the target gene, n=3. i, Fluorescence images of H3K9ac (red) protein in skin tissues of mice of different ages (DAPI, blue), histogram shows the relative quantification of the red signal within the white dashed line in i, n=9. j, Fluorescence images of H3K9ac (red) proteins in skin tissues of one- and two-year old mice after red light irradiation treatment (DAPI, blue). histogram shows the relative quantification of the red signal within the white dashed line in j, n=9. In i and j, only the H3K9ac fluorescence signal within the epidermal region enclosed by the white dotted lines was quantified. k, The evels of H3K9ac and H3 proteins in the skin of mice in groups 2Y and 2Y+R, n=3. l and m, The levels of P16(l) and LAMB1(m) proteins in the skin of mice in groups 4M, 2Y and 2Y+R. n, The expression heatmap of SASP inflammatory factors in the skin of mice in groups 2Y, and 2Y+R, n=3. o, p, q and r, PET/CT images (o) and ^18^F-FDG Radiological signals in skin (p), liver (q) and brain (r) of mice in the 4M, 2Y and 2Y+R treatment groups, n=6. s, Relative content of Acetyl-CoA in the skin tissues of mice in groups 4M, 2Y and 2Y+R, n=5. *P* values were calculated applying an unpaired Student’s *t*-test with unequal variances. ns = not significant (*P* > 0.05).

To further investigate the effects of red light on skin tissue aging in vivo, we established a mouse model of natural aging under conditions of periodic red light irradiation **(Fig S1f** and **g)**. As shown in **Fig S1g**, 2-year-old mice exhibited pronounced facial aging characteristics compared to 4-month-old mice, including significant whisker loss and age-related ocular pathology. During the treatment protocol for our naturally aging mouse model, 1-and 2-year-old mice underwent three and nine 15-day red light irradiation treatments, respectively **(Fig S1f)**. To preclude potential neuromodulatory effects, we employed restraints and light-blocking fabric during red light exposure to restrict the movement of the mice and prevent direct illumination of the eyes **(Fig S1h)**. Prior to and following red light irradiation, we employed infrared detectors to monitor changes in the surface body temperatures of the mice. The results indicated that after exposure to 80 J/cm² of red light, the temperature of the mice increased from 35.9°C to 36.1°C **(Fig S1i)**. This demonstrates that our irradiation protocol did not induce a significant thermos effect. Subsequent analysis of skin and tissue samples from the animal model revealed that periodic red light irradiation did not induce pathological changes in any organ or tissue, including the brain, heart, liver, spleen, lungs, kidneys, or skin **(Fig S1j)**. Indeed, beneficial alterations in aging characteristics were observed across multiple organs and tissues, such as a marked reduction in monocyte numbers within the cardiac muscle tissue **(Fig S1k-q)**. A marked decline in H3K9ac levels was observed in the stratum corneum with advancing age **(Fig 1i)**. In contrast, following periodic red light treatment, H3K9ac levels were increased in the stratum corneum of both 1-year-old and 2-year-old mice **(Fig 1j)**, along with a marked elevation in total H3K9ac within aged skin tissue **(Fig 1k)**. Furthermore, we quantified the relative fluorescence intensity of the P16 protein and the proportion of LAMB1-positive cells in the aged skin tissue. We found that periodic red light treatment not only reduced P16 protein expression and increased the proportion of LAMB1-positive cells (**Fig 1l** and **m)** but also decreased the expression levels of certain SASP inflammatory factors in the skin tissue, such as IL-1β and IL-6 **(Fig 1n)**. This shift in the expression patterns of aging markers and age-related inflammatory factors suggests the anti-aging effects of periodic red light irradiation on the skin.

Another prominent feature of aged tissues is a decline in cellular metabolism. Therefore, we further examined the metabolism of ¹□F-FDG in the skin tissues of aged mice using PET-CT. By capturing radioactive signals, we found that periodic red light irradiation not only increased the radioactive signal in the cortex of aged mice (**Fig 1o** and **p)** but also markedly elevated uptake in other organs, such as the liver and brain **(Fig 1p, q, and r)**. This indicates that red light promotes systemic metabolism in aged mice. Moreover, red light irradiation elevated the acetyl-CoA levels in the skin tissues of aged mice **(Fig 1s)**. As a key intracellular metabolite, acetyl-CoA serves as the sole substrate for histone acetylation and reflects cellular and tissue metabolic activity. These findings suggest that periodic red light treatment may enhance metabolic processes, thereby promoting histone acetylation.

### 2. Red light irradiation increases the metabolic flux and cellular acetyl-CoA content in keratinocytes

To identify the metabolic remodeling effects of red light, building upon the promotion of keratinocyte activity observed at 80 J/cm² **(Fig 1a)**, we further examined the effects of red light on cell migration and proliferation. Scratch assay results demonstrated that the wound closure rate in keratinocytes increased from 40% to 60% 24 h after red light treatment **(Fig 2a)**. Moreover, within 24 h of red light irradiation, the keratinocyte proliferation rates were markedly higher than those in the control group **(Fig 2b)**. Since cellular metabolic levels are closely linked to proliferation, viability, and migration, we further assessed keratinocyte uptake of two major metabolic substrates—glucose and fatty acids—from the culture medium within 12 h of red light treatment. As demonstrated in **Fig 2c** and **Fig 2d**, after 12 h of red light treatment, the uptake of both glucose and fatty acids from the medium increased by approximately 10% compared to that in the control group. Under this conditions, the mitochondrial membrane potential was significantly increased **(Fig 2e)**. These findings indicate that red light irradiation enhances the metabolic flux in keratinocytes and boosts mitochondrial metabolic function.

**Figure 2.**
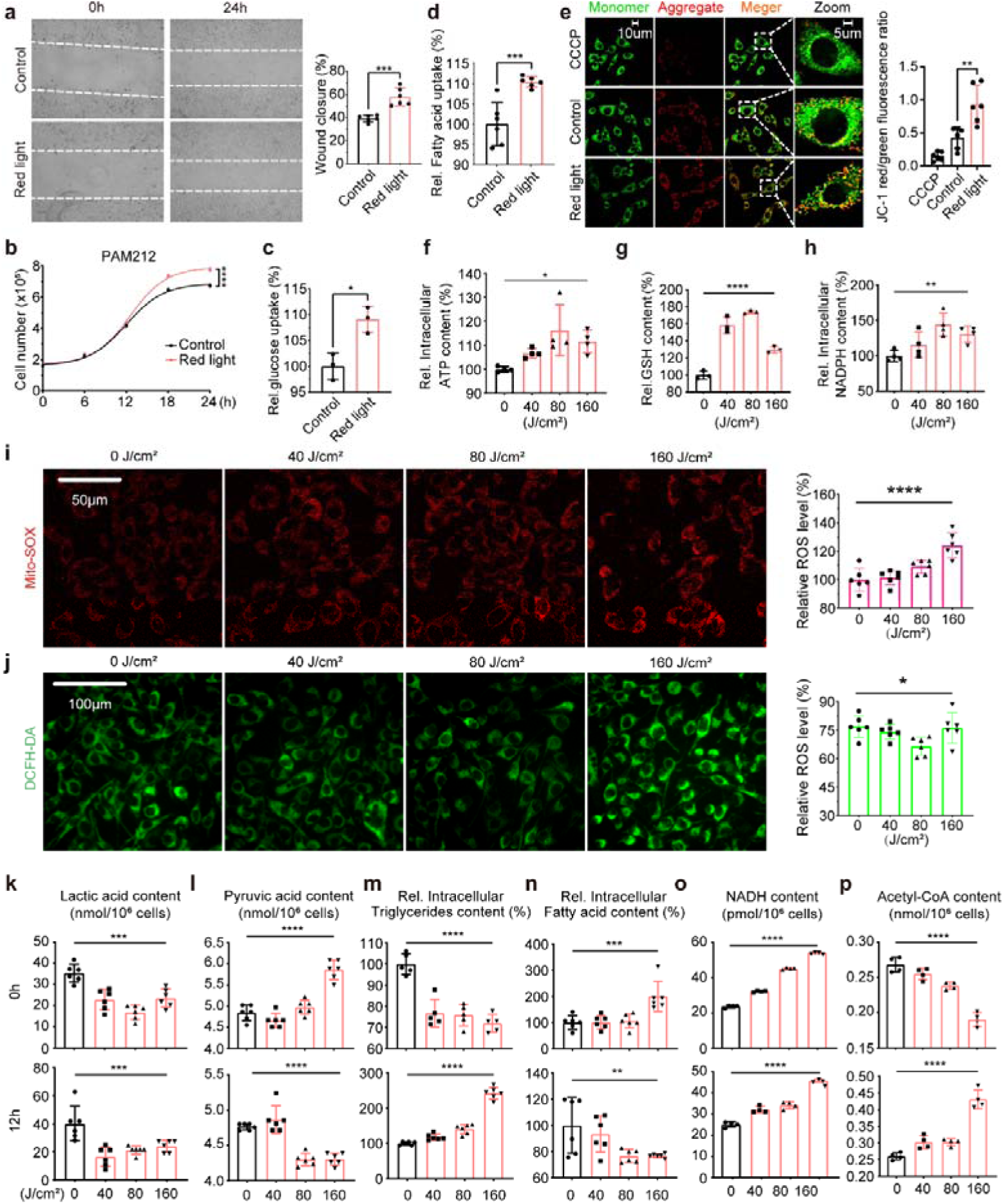
Red light irradiation promotes keratinocyte metabolic reprogramming. **a,** Migration of keratinocytes after 24 h of red light treatment (n = 6). **b,** Proliferation of keratinocytes: 0–24 h after red light treatment (n = 3). **c** and **d,** Uptake of glucose (**c**, n=3) and fatty acid (**d**, n=6) in the culture medium by keratinocytes 24 h after red light treatment. **e,** Mitochondrial membrane potential of keratinocytes after red-light treatment. CCCP, Carbonyl cyanide m-chlorophenylhydrazone (n=6). **f,** Graded-dose red light treatment affects ATP production in keratinocytes (n = 4). **g** and **h,** Relative changes in cellular GSH (**g**) and NADPH (**h**) contents 12 h after treatment of keratinocytes with graded doses of red light (n=3). **g** and **h,** Fluorescence images of total reactive oxygen species (ROS) levels (**i**, n=6) and mitochondrial ROS levels (**j**, n=6) in keratinocytes under different red light doses. **k-p,** Relative changes in intracellular lactate (**k**, n=6), pyruvate (**l**, n=6), triglyceride (**m**, n=6), fatty acid (**n**, n=6), NADH (**o**, n=4), and acetyl-CoA (**p**, n=4) at 0h and 12h after treatment of keratinocytes with graded doses of red light. ANOVA was used to analyse the significant differences in **f-p**. *P* values were calculated applying an unpaired Student’s *t*-test with unequal variances. ns = not significant (*P* > 0.05).

Interestingly, upon further examination of key metabolites associated with intracellular redox status and glycolytic pathways across a red light dose gradient of 0–160 J/cm^2^, we observed a pronounced biphasic dose response in ATP levels and the two primary reducing agents, GSH and NADPH **(Fig 2f, g**, and **h)**. The concentrations of ATP, GSH, and NADPH increased with red light irradiation dose within the 0–80 J/cm² range, revealing a potential reason for keratinocyte activity peaking at 80 J/cm² red light conditions and indicating that red light promotes cellular metabolism **(Fig 1a** and **Fig 2f-h)**. Moreover, our assessment of the mitochondrial membrane potential and cellular reactive oxygen species (ROS) in keratinocytes under varying red light doses revealed that mitochondrial ROS production increased with increasing red light exposure **(Fig 2i)**. Interestingly, the overall level of reactive oxygen species within keratinocytes exhibited a decreasing trend across the red light dose range of 0–80 J/cm² **(Fig 2j)**. Considering that cytoplasmic GSH and NADPH levels also increased with increasing red light doses within the 0–80 J/cm² range (**Fig 2g** and **h)**, we propose that the biphasic dose effect driven by red light is most likely mediated through the neutralization of mitochondrial ROS by cellular reducing agents. Excessively high red light doses induce excessive mitochondrial ROS production, impair normal cellular processes, and consequently elicit adverse reactions. This is further supported by the observation that the mitochondrial membrane potential levels in keratinocytes begin to decline from their peak at 160 J/cm² red light exposure **(Fig S2a)**.

Regarding changes in metabolite levels following red light irradiation, we compared intracellular lactate, pyruvate, triglyceride, and fatty acid concentrations at 0 and 12 h post-red light exposure **(Fig 2k-n)**. We observed that, apart from lactate exhibiting a sustained decline within 12 h post- red light, the other three metabolites demonstrated distinct temporal patterns of change within 12 h of red light treatment **(Fig 2k-n)**, with increased cellular utilization of fatty acids (**Fig 2m** and **n)**. These findings suggest that the metabolic effects of red light involve accelerated aerobic metabolism and fatty acid oxidation. Based on the conversion relationships between the aforementioned metabolic pathways, as depicted in **Fig 1s**, we reassessed the NADH levels in keratinocytes at 0 and 12 h after red light treatment. The results revealed an increasing trend in NADH content at both time points **(Fig 2o)**. These findings conclusively demonstrate that red light irradiation promotes cellular metabolic processes, including glycolysis, fatty acid metabolism, and the TCA cycle. This aligns with the observations in animal aging models. Finally, we measured the acetyl-CoA levels in keratinocytes following red light irradiation. Although red light rapidly depleted acetyl-CoA, both acetyl-CoA content and H3K9ac levels increased with increasing red light dose at 12 h post-treatment (**Fig 2p** and **1b)**. The concentration of acetyl-CoA in the cell nucleus also increased following exposure to red light (**Fig S2b**). These results further demonstrates the potential of red light to influence cellular aging by driving histone acetylation through metabolic regulation.

### 3. Red light modulates cellular metabolism by activating the PI3K-AKT and PPAR-α metabolic pathway

Red light enhanced cellular metabolic flux, thereby promoting acetyl-CoA accumulation in keratinocytes and aged skin tissue, which may contribute to increased histone acetylation and aging mitigation (**Fig. 3a**). To further investigate how red light regulates glycolipid metabolism, we performed transcriptomic analysis of keratinocytes after red-light treatment. KEGG enrichment analysis showed that the top 20 significantly enriched pathways included several metabolism- and aging-related pathways, such as the p53 and MAPK signalling pathways (**Fig. 3b**). Consistently, differentially expressed genes showed an overall trend toward upregulation (**Fig. 3c**), suggesting that red light broadly enhances transcriptional activity. Given the close association between red-light-mediated photobiological effects and cellular metabolism, we focused on metabolically relevant pathways, particularly the PI3K-AKT and PPAR signalling pathways, which are involved in glucose metabolism and fatty acid metabolic regulation, respectively (**Fig. 3b**). Further analysis revealed that red light upregulated multiple metabolism-related genes, including genes involved in glycolysis, the tricarboxylic acid cycle, the pentose phosphate pathway, and fatty acid metabolism. These included the fatty acid transporter gene *Cpt1a*, the key FAO-related transcription factor *Ppara*, and several PI3K family genes (**Fig. 3d** and **Fig. S3a**). Fatty acid metabolism and glycolysis/gluconeogenesis pathways were enriched, with the upregulation of fatty acid metabolism being the most pronounced across all pathways **(Fig S3b)**. These findings suggest that red light may activate cellular glycolytic and lipid metabolism by separately stimulating the aforementioned signalling pathways, thereby mobilizing overall cellular metabolic vigor.

**Figure 3.**
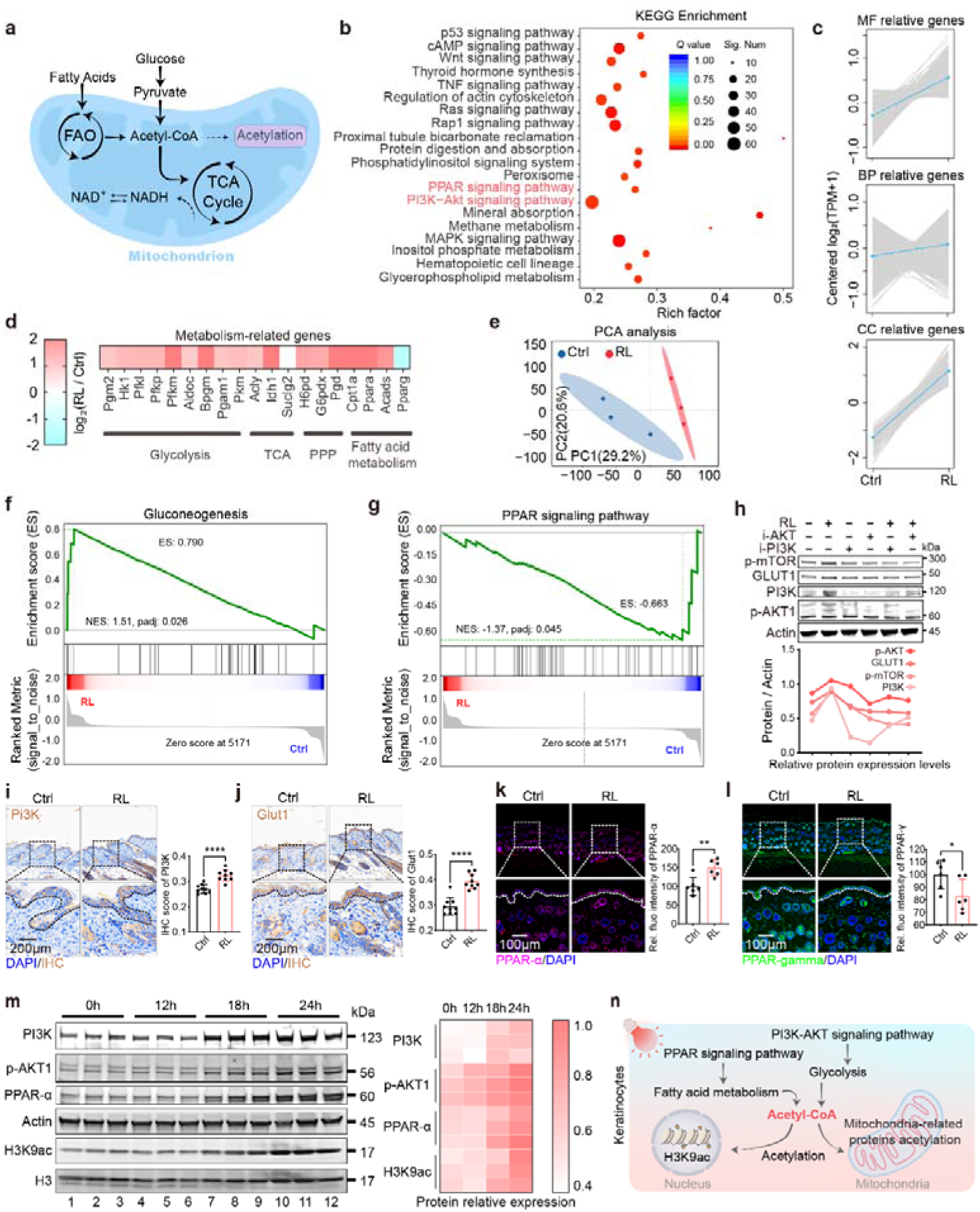
Red light activated the PI3K-akt signalling pathway and PPAR-α in keratinocytes simultaneously with H3k9ac. **a,** Schematic representation of substance metabolism in mitochondria. **b,** Scatterplot of significantly enriched functions for transcript data from keratinocytes treated with red light (Top 20 KEGG for plotting). **c,** Differential gene module expression trend line graph. Coloured lines indicate the mean value of the change for this group of genes. **d,** Expression of keratinocyte metabolism-related genes after red light treatment. **e,** Principal component analysis (PCA) of keratinocyte proteomics in the red light and dark treatment groups, n=3. **f** and **g,** GSEA analysis of gluconeogenesis (**f**) and PPAR signalling pathway (**g**) proteins in keratinocytes after red light irradiation. **h,** Expression levels of p-mTOR, Glut1, PI3K, and p-AKT proteins in keratinocytes after red light treatment in the presence of AKT and PI3K inhibitors (n = 3). **i** and **j,** Immunohistochemical staining of PI3K (**i**) and Glut1(**j**) proteins in skin tissues after red light irradiation, n=9. **k** and **l,** Immunofluorescence staining of PPAR-α (**k**) and PPAR-γ (**l**) proteins in skin tissues after red light irradiation (n = 6). **m,** Changes in the levels of PI3K, p-Akt, PPAR-α, Actin, H3K9ac and H3 proteins in keratinocytes at 0h, 12h, 18h, and 24h after red light treatment, n=3. **n,** Schematic diagram of increased glycolipid metabolism driving histone acetylation in cells after red light treatment of keratinocytes. Ctrl: Control; RL: Red light. *P* values were calculated applying an unpaired Student’s *t*-test with unequal variances. ns = not significant (*P* > 0.05).

Similarly, proteomic analysis of keratinocytes following red light treatment revealed that both dark and red light conditions significantly influenced protein expression patterns within keratinocytes **(Fig 3e)**. Genome-wide signature analysis (GSEA) of differentially expressed proteins further enriched the gluconeogenesis and PPAR signalling pathways in keratinocytes under dark versus red light conditions (**Fig 3f** and **g)**. Within the proteomic findings, we further observed elevated levels of proteins belonging to the GPCRs family **(Fig S3c** and **d)**. As upstream regulators of the PI3K-AKT signalling pathway, the enrichment of molecular processes involving GPCRs in the proteomic analysis of keratinocytes following red light exposure indirectly indicates the regulatory role of red light in the PI3K-AKT pathway.

In summary, these findings suggest that red light treatment activates intracellular glycolytic and lipid metabolic pathways. Consequently, we validated these two signalling pathways at both the cellular and tissue levels. First, we examined the effects of red light on glycolytic pathways. Given that cellular glycolysis is primarily regulated through the coordinated action of the metabolic control centre mTOR and the PI3K-AKT signalling pathway, we initially assessed changes in key protein levels within these pathways at the cellular level. We observed a marked increase in protein levels along the mTOR/PI3K-AKT/Glut1 signalling axis following red light treatment **(Fig 3h)**. Inhibition of this axis was achieved by administering either a PI3K inhibitor or an AKT inhibitor. Crucially, red light treatment under inhibitor conditions partially mitigated this suppression, confirming the role of red light in promoting protein expression along this axis at the cellular level **(Fig 3h)**. At the tissue level, we validated the expression of PI3K and Glut1 in the skin tissue following red light treatment. Immunohistochemical staining results also revealed increased PI3K and Glut1 signalling (**Fig 3i** and **j)**.

Furthermore, we independently validated alterations in PPAR-α and PPAR-γ levels, which are key regulators of lipid metabolism and synthesis, at both cellular and tissue scales. After red light treatment, cellular and tissue PPAR-α levels increased, whereas PPAR-γ levels decreased (**Fig 3k** and **l)**. To characterize the regulation of H3K9ac by red light-activated glycolipid metabolism, we examined the temporal dynamics of glycolipid metabolism activation and H3K9ac changes in keratinocytes within 24 h of red light exposure. Surprisingly, the results indicated that key proteins within the PPAR-α/PI3K-AKT signalling pathway began to significantly increase 12 h after red light cessation, occurring almost simultaneously with the increase in H3K9ac levels **(Fig 3m** and **Fig S3e)**. Fluorescence imaging of H3K9ac within 24 h of red light irradiation similarly demonstrated a progressive elevation in H3K9ac levels **(Fig S3f)**. These findings confirm that red light can independently activate the PI3K-Akt and PPAR-α pathways to regulate cellular glycolipid metabolism at the cellular level. Combined with the experimental findings of increased acetyl-CoA levels in keratinocytes at 12 h post-red light irradiation **(Fig 2p)**, we ultimately elucidated a potential mechanism by which red light promotes acetylation-related processes in keratinocytes by activating glycolipid metabolic pathways, leading to cellular acetyl-CoA accumulation **(Fig 3n)**.

### 4. Red light mediates histone acetylation primarily via lipid metabolism

We demonstrated that red light activates intracellular glycolipid metabolic pathways and histone acetylation. However, the mechanism by which mitochondrial metabolism is activated under red light conditions and the relationship between metabolic pathway activation and histone acetylation remain unclear. We need to further clarify whether histone acetylation driven by metabolic activation in keratinocytes under red light irradiation exhibits metabolic specificity and how red light directly influences mitochondrial metabolic processes.

To investigate the direct effects of metabolism on histone acetylation, we first treated keratinocytes with agonists and inhibitors of the key metabolism-related proteins AKT1 and PPAR-α, respectively. Subsequently, we examined the impact of histone acetylation on glycolytic and lipolytic pathways under red light conditions by inhibiting acetyltransferases and deacetylases. Finally, we used inhibitors of glucose and lipid metabolism to establish the regulatory role of red light-induced metabolic activation of H3K9ac under red light exposure. The results revealed that the AKT agonist SC-79 significantly promoted H3K9ac levels (**Fig 4a**, **b** and **Fig S4a)**. However, while the AKT inhibitor MK2206 markedly suppressed p-mTOR levels, it did not significantly inhibit histone acetylation (**Fig 4c**, **d** and **Fig S4b)**. Conversely, intracellular acetyl-CoA levels were markedly increased and decreased by SC-79 and MK2206, respectively (**Fig 4e** and **f)**. Similarly, we performed experiments with the PPAR-α agonist, WY14643, and the PPAR-α antagonist, GW6471. We found that activating PPAR-α markedly promoted H3K9ac, but had no significant effect on p-mTOR or p-AKT levels **(Fig 4g, h** and **Fig S4c)**. Inhibition of PPAR-α similarly reduced histone acetylation **(Fig 4i, j** and **Fig S4d)**. Intracellular acetyl-CoA levels exhibited a pronounced dose-dependent response to both WY14643 and GW6471 (**Fig 4k** and **l)**. These findings conclusively demonstrate that acetyl-CoA derived from lipid metabolism serves as the primary acetyl donor for red-light-driven H3K9ac.

**Figure 4.**
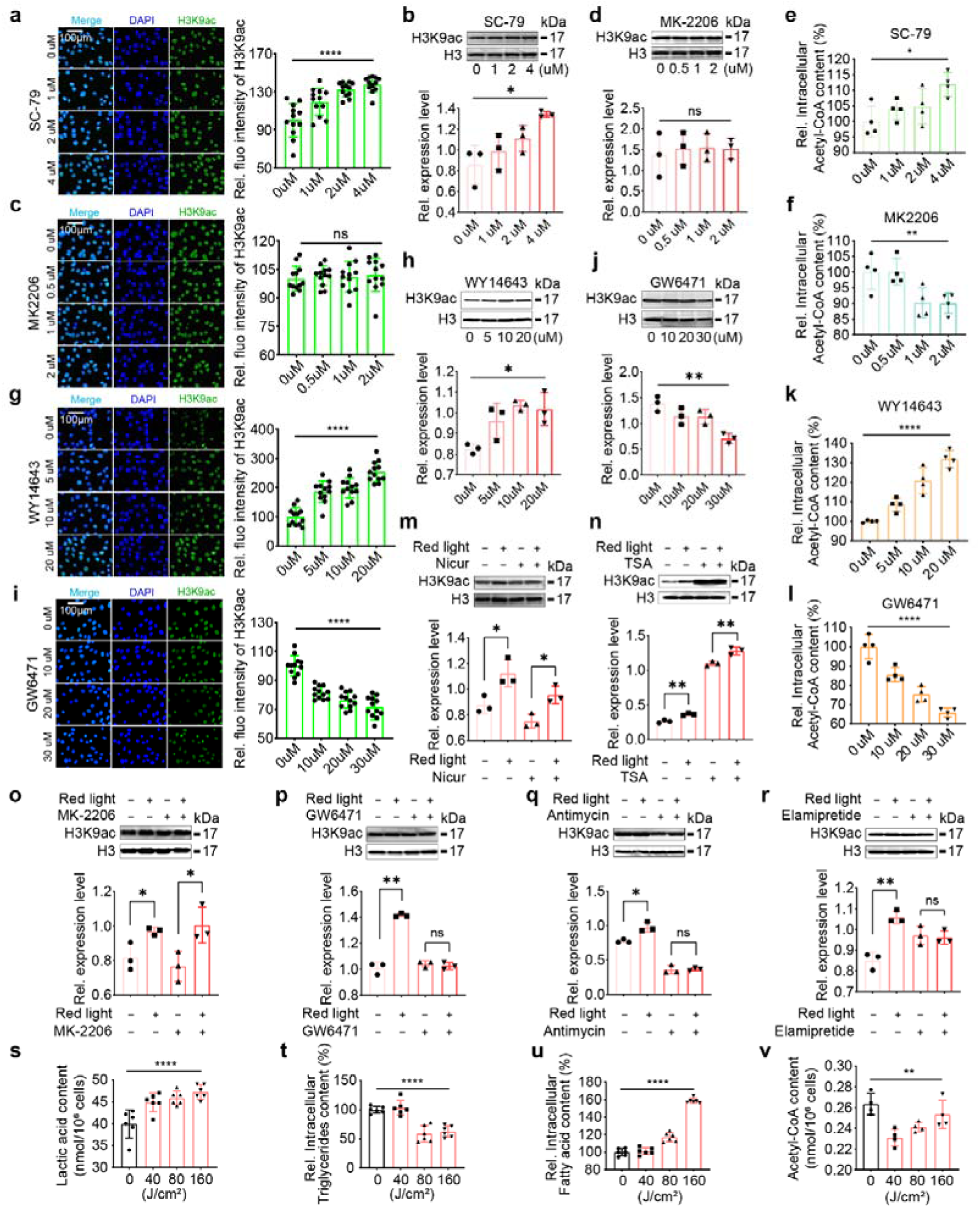
Red light drives metabolism through mitochondria and mediates histone-acetylation primarily through lipid metabolism. **a** and **b,** Fluorescence images of H3K9ac (green, **a**, n=12) protein and levels of H3K9ac and H3 proteins **(b**, n=3) in keratinocytes under different concentrations of SC-79 treatment conditions (DAPI, blue). **c** and **d,** Fluorescence images of H3K9ac (green, **c**, n=12) protein and levels of H3K9ac and H3 proteins **(d**, n=3) in keratinocytes under different concentrations of MK2206 treatment conditions (DAPI, blue). **e** and **f,** Effects of different concentrations of SC-79 **(e)** and MK2206 **(f)** on Acetyl-CoA content in keratinocytes, n=4. **g** and **h,** Fluorescence images of H3K9ac (green, **g**, n=12) protein and levels of H3K9ac and H3 proteins **(h**, n=3) in keratinocytes under different concentrations of WY14643 treatment conditions (DAPI, blue). **i** and **j,** Fluorescence images of H3K9ac (green, **i**, n=12) protein and levels of H3K9ac and H3 proteins **(j**, n=3) in keratinocytes under different concentrations of GW6471 treatment conditions (DAPI, blue). **k** and **l,** Effects of different concentrations of WY14643 **(k)** and GW6471 **(l)** on Acetyl-CoA content in keratinocytes, n=4. **m** and **n,** Effect of red light treatment on the levels of H3K9ac and H3 proteins in keratinocytes in the presence of 5uM Nicur **(m)** and TSA **(n)**, n=3**. o** and **p,** Effect of red light treatment on the levels of H3K9ac and H3 proteins in keratinocytes in the presence of 2uM Mk2206 **(o)** and 20uM GW6471 **(p)**, n=3. **q** and **r,** Effect of red light treatment on the levels of H3K9ac and H3 proteins in keratinocytes in the presence of 5uM Antimycin **(q)** and Elamipretide **(r)**, n=3. **s, t, u,** and **v,** Relative contents of lactate **(s**, n=6), triglycerides **(t**, n=6), fatty acids **(u**, n=6) and Acetyl-CoA **(v**, n=4) in keratinocytes after irradiation of 5uM Elamipretide-treated keratinocytes with graded doses of red light. *P* values were calculated applying an unpaired Student’s *t*-test with unequal variances. ns = not significant (*P* > 0.05).

In subsequent experiments, we inhibited histone acetylation using the histone acetyltransferase inhibitor Nicur (a curcumin-derived p300/CBP inhibitor) and observed that Nicur appeared to exert no discernible effect on the glycolytic pathway. Moreover, following red light treatment, both the intracellular PI3K/Akt/mTOR metabolism-related signaling pathway and H3K9ac exhibited pronounced activation trends **(Fig 4m** and **Fig S4e)**. Conversely, when histone acetylation was inhibited using the deacetylases inhibitor Trichostatin A (TSA), the levels of proteins associated with the PI3K/Akt/mTOR metabolism-related signaling pathway showed a decreasing trend **(Fig 4n** and **Fig S4f)**. In contrast, TSA treatment markedly increased PPAR-α expression, and red light irradiation further amplified this effect **(Fig S4f)**. This regulatory effect was not pronounced under Nicur treatment **(Fig S4e)**. In both sets of experiments, red light treatment increased intracellular H3K9ac levels **(Fig 4m, n** and **Fig S4g, h)**. These findings further validate that red light-induced metabolic activation directly causes H3K9ac increase rather than indirectly through regulation of histone acetyltransferases and histone deacetylases. These results indicate that histone acetylation preferentially promotes the expression of lipid metabolism-related proteins. Further treatment of keratinocytes with the AKT1 inhibitor MK2206 revealed that red light still upregulated H3K9ac, whereas treatment with the lipid metabolism inhibitor GW6471 abolished the ability of red light to increase intracellular H3K9ac levels(**Fig 4o** and **p)**. This demonstrates that red light-driven lipid metabolism plays a fundamental role in elevating H3K9ac levels **(Fig 4o, p** and **Fig S4i, j)**.

Existing research indicates that cytochrome c oxidase is the direct target of red light within mitochondria^48^. We propose that the transient effect of red light on cellular metabolic regulation (changes in intracellular metabolite levels at 0 h after red light exposure) is also linked to the mitochondrial metabolic activity **(Fig 2k-p)**. Consequently, we independently inhibited the electron transport chain and cytochrome c oxidase in keratinocytes treated with red light. The results revealed that disruption of the electron transport chain abolished the regulatory capacity of red light over metabolic pathways and histone acetylation **(Fig 4q** and **Fig S4k)**. Similarly, under cytochrome c oxidase inhibition, red light treatment failed to increase histone acetylation and consequently lost its regulatory effect on the metabolic pathway **(Fig 4r** and **Fig S4l)**. Furthermore, following cytochrome c oxidase inhibition, the pattern of transient intracellular key metabolite regulation after red light irradiation changed **(Fig 4s-v)**, and the effects of red light on lactate and acetyl-CoA content were reversed **(Fig 2k, p** and **Fig 4s, v)**. Additionally, cellular component enrichment analysis of the acetylated proteomics data also revealed that multiple mitochondrial processes were significantly upregulated following red light treatment **(Fig S4c** and **d)**. Collectively, these findings demonstrate that red light-induced transient metabolic drive is closely linked to mitochondrial function.

Based on these experimental findings, we elucidated the mechanism by which red-light irradiation modulates intracellular H3K9ac levels. Specifically, red light drives mitochondrial metabolism in keratinocytes, altering metabolite concentrations and activating relevant signalling pathways. This ultimately manifests the characteristic biological regulatory effect of red light: red light activates intracellular glycolipid metabolism and primarily promotes histone acetylation within keratinocytes via acetyl-CoA supplied by fatty acid metabolism.

### 5. Downregulation of SIRT4 induced by red light alleviates oxidative aging in keratinocytes

Although red light exerts transient regulatory effects by directly driving mitochondrial metabolism, the mechanisms and molecular targets by which it subsequently activates PPAR-α related fatty acid metabolism pathway and mediates H3K9ac remain unclear.

Among the changes in keratinocyte total protein expression, the significant reduction in SIRT4 protein levels following red light treatment drew our attention (**Fig 5a** and **b)**. As a member of the Sirtuin protein family, SIRT4 possesses deacetylase activity. The reduction in SIRT4 levels leads to increased protein acetylation, consistent with our experimental findings. Furthermore, as a deacetylase primarily distributed in the mitochondria, SIRT4 has been reported to regulate cellular metabolism, particularly fatty acid oxidation processes, by modulating the acetylation of mitochondrial metabolism-related proteins **(Fig 5c)**. Moreover, SIRT4 modulates PPAR-α activation by inhibiting SIRT1 activity, thereby regulating NF-κB-mediated inflammatory signalling^49^. This aligns with our observation that red light irradiation remodels inflammatory factor expression in oxidatively aged keratinocytes and tissues, as well as in naturally aged skin **(Fig S1e** and **Fig 1n)**. Therefore, we propose that the reduction in SIRT4 protein levels following red light irradiation may be the primary mechanism underlying the enhanced mitochondrial metabolism observed after red light treatment.

**Figure 5.**
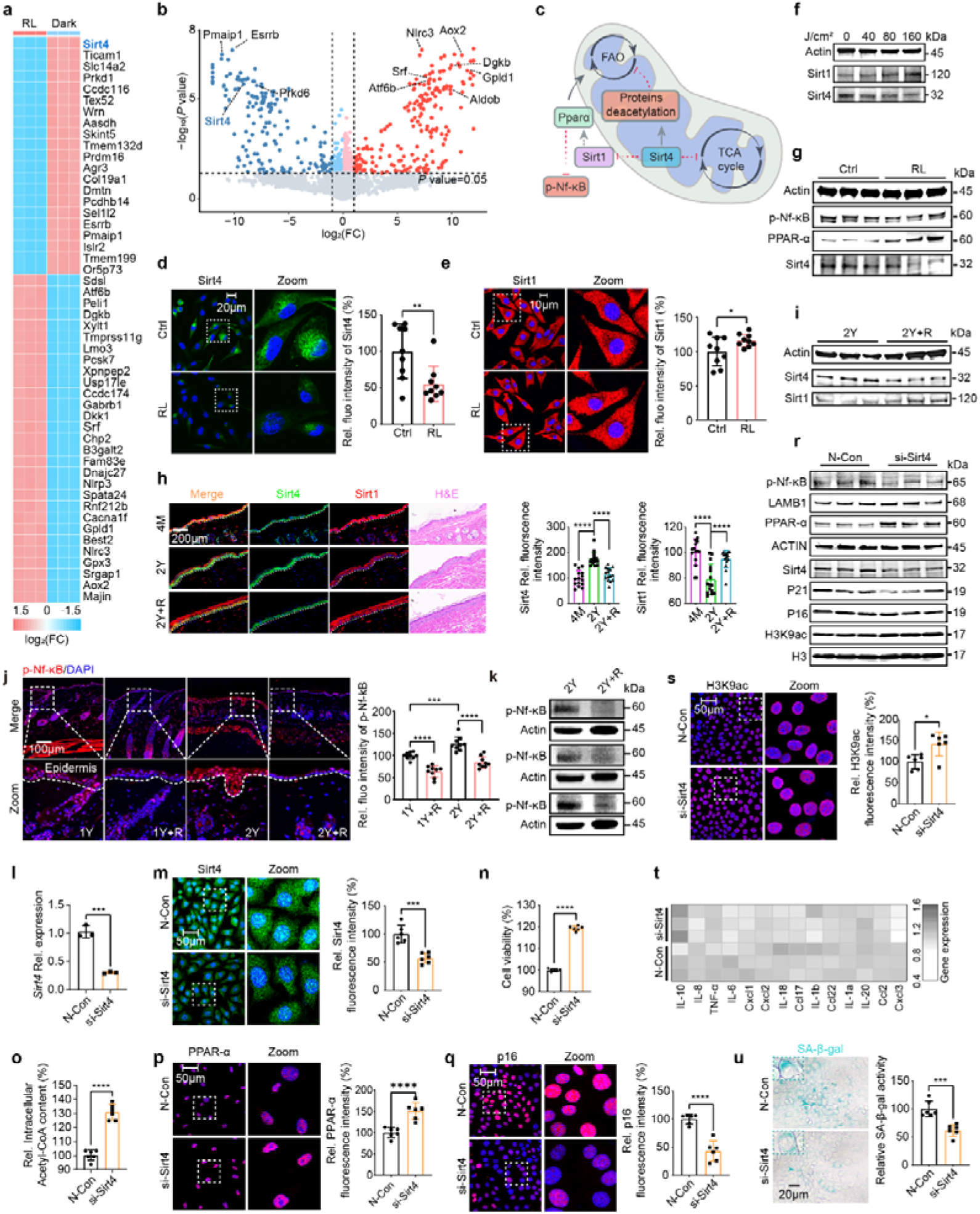
Red light-induced downregulation of SIRT4 is primarily responsible for the activation of lipid metabolism, alleviation of inflammation, and resistance to aging in keratinocytes. **a**, Heatmap showing the top 50 significantly differentially expressed proteins in red light-irradiated keratinocytes. **b,** Volcano plot of protein content changes in keratinocytes after red light treatment. **c,** Schematic representation of the role of SIRT4 in mitochondria and cell. **d** and **e,** Fluorescence images of SIRT4 (**d**) and SIRT1 (**e**) protein and nucleus in keratinocytes after red light treatment, n=9. **f,** Intracellular protein levels of SIRT1 and SIRT4 after dose-gradient red light treatment of keratinocytes, n=3. **g,** Protein levels of p-Nf-κB, PPAR-α and SIRT4 in keratinocytes after red light treatment, n=3. **h,** H&E staining images and fluorescence images of SIRT4 (green) and SIRT1 (red) proteins of mouse skin tissues in 4M, 2Y and 2Y+R treatment groups (DAPI, blue), n=3. **i,** The levels of SIRT4 and SIRT1 proteins in the skin of aged mice after cyclic red light treatment, n=3. **j** and **k,** Fluorescence images of p-Nf-κB (red, **j**) in mice skin tissues after red light treatment of 1Y and 2Y senescent mice (DAPI, blue) and the levels of p-Nf-κB protein (**k**) in skin tissues of 2Y senescent mice after red light treatment, n=3. **l** and **m,** Relative expression of SIRT4 mRNA (**l**, n=3) and fluorescence images of SIRT4 protein (green) (**m**, n=6) in keratinocytes after silencing of the SIRT4 gene in keratinocytes using siRNA. **n** and **o,** Changes in cellular activity (**n**, n=3), relative content of intracellular acetyl-CoA (**o**, n=6) in keratinocytes after silencing the SIRT4 gene in keratinocytes using siRNA. **p** and **q,** Fluorescence images of PPAR-α (**p**) and P16 (**q**) proteins in keratinocytes after silencing the SIRT4 gene in keratinocytes using siRNA, n=6. **r,** Protein levels of p-Nf-κB, LAMB1, PPAR-α, SIRT4, p21, p16, H3K9ac, and H3 in keratinocytes after silencing the SIRT4 gene in keratinocytes using siRNA, n=3. **s,** Fluorescence images of H3K9ac proteins in keratinocytes after silencing the SIRT4 gene in keratinocytes using siRNA, n=6. **t,** Relative expression of SASP inflammatory factors in keratinocytes following silencing of the SIRT4 gene in keratinocytes using siRNA, n=3. **u,** Images of β-galactosidase staining in keratinocytes after silencing the SIRT4 gene in keratinocytes cells using siRNA, n=6. N-Con: Negative Control; RL: Red light. *P* values were calculated applying an unpaired Student’s *t*-test with unequal variances. ns = not significant (*P* > 0.05).

To further validate the proteomics analysis results and our hypothesis, we first verified SIRT4 expression in keratinocytes exposed to red light. We found that SIRT4 levels progressively decreased with increasing light doses (**Fig 5d** and **f)**, whereas SIRT1 levels increased (**Fig 5e** and **f)**. Moreover, under red light treatment, the trend in SIRT4 levels mirrored that of p-NF-κB, while exhibiting the opposite pattern to that of PPAR-α **(Fig 5g)**. These findings validate the regulatory capacity of SIRT4 on intracellular lipid metabolism targets and inflammatory signalling pathways. We also assessed SIRT1/4 protein expression levels in aged animal models. Similarly, we observed a progressive increase in SIRT4 expression in tissues with advancing age, whereas SIRT1 expression exhibited an inverse trend (**Fig 5h** and **i)**. Red light treatment significantly downregulated SIRT4 protein expression in aged skin tissue (**Fig 5h** and **i)**. Moreover, age-related p-NF-κB activation in the skin tissue was markedly suppressed following periodic red light exposure **(Fig 5j, k** and **Fig S5a).** At both the cellular and tissue levels, we observed a negative correlation between PPAR-α and p-NF-κB protein expression levels **(Fig S5b, c)**. This negative correlation was more pronounced in keratinocytes under red light **(Fig S5b)**. These findings suggest that reduced SIRT4 protein content plays a substantial role in the red light-driven cellular anti-aging effects.

Therefore, to further elucidate the role of SIRT4 in cellular senescence, we employed siRNA to silence SIRT4 expression in keratinocytes. Following a marked reduction in both transcriptional and protein expression levels of SIRT4 within the keratinocytes (**Fig 5l** and **m)**, we observed a significant increase in cellular activity, along with elevated intracellular acetyl-CoA levels (**Fig 5n** and **o)**. These findings demonstrate that SIRT4 downregulation promotes cellular metabolism. Furthermore, we observed a marked increase in PPAR-α protein levels, activation at histone H3K9 sites, reduced expression of senescence factors p16 and p21, and elevated Lamin-B1 in SIRT4-silenced cells **(Fig 5p-s)**. Nf-κB phosphorylation levels and SASP-associated inflammatory cytokine secretion in the si-SIRT4-treated group exhibited a similar expression remodelling to that observed in the red light treatment group (**Fig 5r** and **t)**. Furthermore, by measuring galactosidase activity in keratinocytes following si-SIRT4 treatment, we found that SIRT4 downregulation significantly alleviated senescence in the keratinocyte oxidative aging model **(Fig 5u)**. These findings conclusively demonstrate the positive role of SIRT4 in keratinocyte anti-aging processes, revealing that red light-driven downregulation of SIRT4 protein levels may constitute a key mechanism underlying these effects.

### 6. Red light activates fatty acid oxidation by inhibiting the SIRT4-MCD axis

Given that SIRT4 is a mitochondrial-localised deacetylase, current research suggests that SIRT4 participates in the regulation of mitochondrial fatty acid oxidation by deacetylating malonyl-CoA decarboxylase (MCD)^50^. This is consistent with our observation that red light mediates H3K9ac via lipid metabolism. Therefore, in order to further investigate and elucidate the mechanisms underlying the anti-aging effects of red light, we conducted acetylation proteomic analysis of keratinocytes following red light treatment.

Our analysis revealed increased acetylation levels in numerous metabolism-related proteins after red light irradiation. These proteins are involved in metabolic processes linked to mitochondrial function, such as citrate synthase (CS) and MCD (**Fig 6a** and **b)**, consistent with the mitochondrial localization and function of SIRT4 (**Fig 5c**). Furthermore, acetylated proteomics data were significantly enriched in multiple molecular processes involving acetyl-CoA oxidase/dehydrogenase and histone-related processes **(Fig 6c** and **Fig S6a, b)**, alongside increased acetylation of mitochondrial electron transport chain complex proteins **(Fig S6c).** Proteins showing increased acetylation following red light irradiation were implicated in biological processes, including fatty acid catabolism, oxidative phosphorylation, the TCA cycle, and pyruvate metabolism **(Fig 6d)**. KEGG pathway analysis of acetylated proteins revealed significant enrichment in fatty acid catabolism and the TCA cycle **(Fig 6e)**. GO analysis identified significantly enriched biological processes, including fatty acid oxidation, histone acetyltransferase activity, and histone deacetylation **(Fig S6d** and **e)**. Furthermore, combined proteomics and acetylated proteomics analyses revealed that red light-promoted molecular processes were associated with fatty acid oxidation, the TCA cycle, acetyl-CoA transfer, and histone deacetylation processes **(Fig 6f)**. The integration of these multi-omics analyses with existing experimental validation fully elucidates the role of red light in activating intracellular fatty acid oxidation and mitochondrial metabolism.

**Figure 6.**
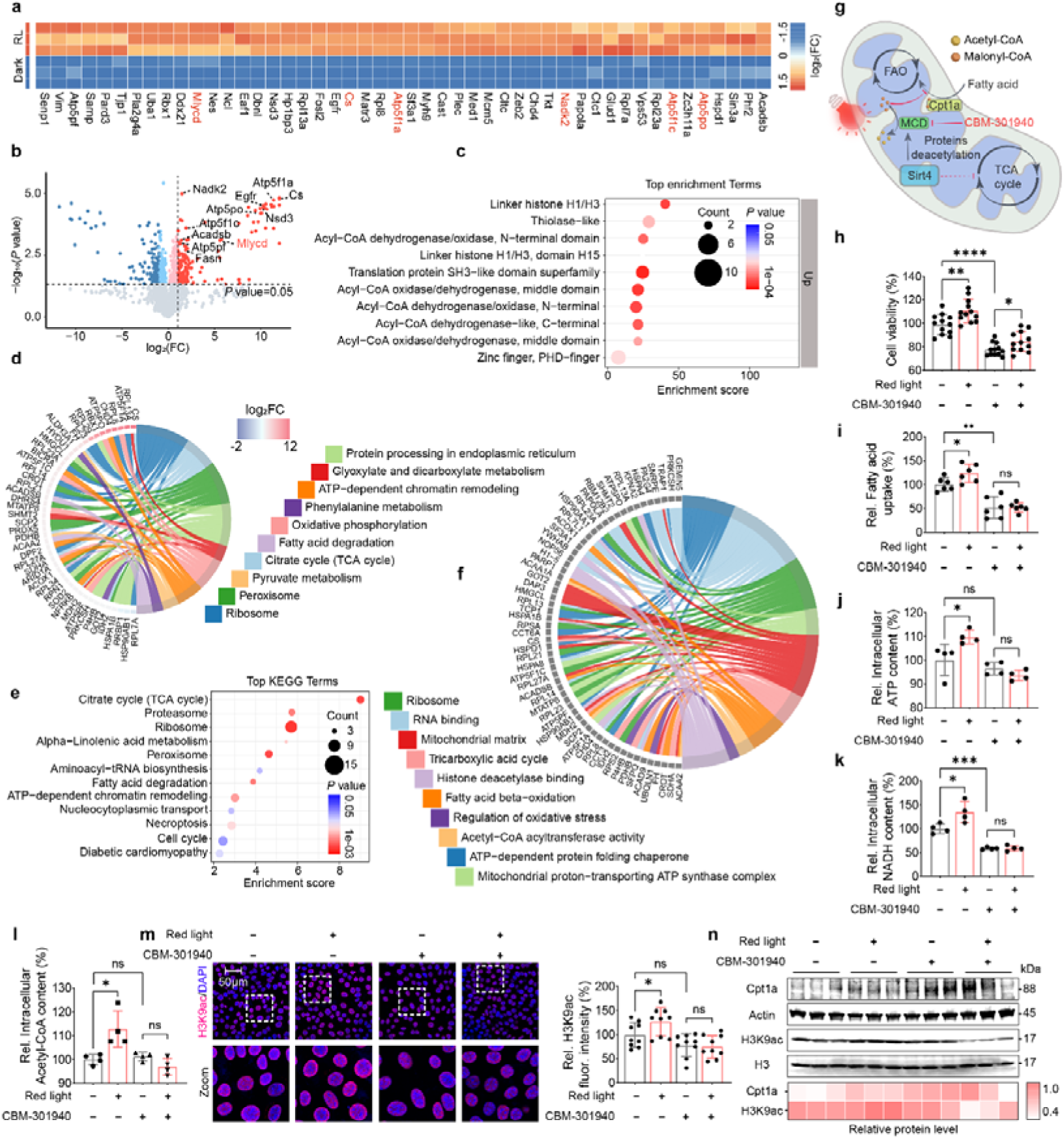
Red light activates lipid metabolism and increase H3K9ac levels in keratinocytes via SIRT4 downregulation-mediated MCD acetylation. **a**, Heatmap showing the top 50 significantly differentially expressed acetylated proteins in red light-irradiated keratinocytes. **b,** Volcano plot of changes in the content of acetylation-modified proteins in keratinocytes after red light treatment. **c,** Acetylation modification proteomics top10 protein functional enrichment analysis (Protein functional annotation by InterPro). **d,** Proteomic top10 enriched pathway chordal map of red light irradiated keratinocytes (KEGG analysis). **e,** Top12 molecular process enriched by combined analysis of keratinocyte proteomics and acetylation modification proteomics after red light treatment (KEGG analysis). **f,** Top 10 pathways chordal map enriched by combined analysis of keratinocyte proteomics and acetylation modification proteomics after red light treatment. **g,** Schematic representation of the role of SIRT4 in mitochondria FAO and TCA cycle progress. **h, i, j, k** and **l,** Effects of red light on cell viability (CCK-8, **h**, n=12), fatty acid uptake (**i**, n=6), ATP production (**j**, n=4), NADH levels (**k**, n=4), and Acetyl-CoA content (**l**, n=4) in keratinocytes treated with 5 μM CBM-301940. **m** and **n,** Representative immunofluorescence staining of H3K9ac (**m**, n=9)and Western blot analysis of CPT1A, β-Actin, H3K9ac and H3 in keratinocytes treated with red light and 5 μM CBM-301940 (**n**, n=3). *P* values were calculated applying an unpaired Student’s *t*-test with unequal variances. ns = not significant (*P* > 0.05).

To further explore the direct link between red light and lipid metabolism, we validated the effects of the SIRT4-MCD axis on cellular fatty acid oxidation and H3K9ac at the keratinocyte level under red light conditions. Within mitochondria, SIRT4 directly inhibits MCD activity through deacetylation, preventing the conversion of malonyl-CoA to acetyl-CoA^50, 51^. The resulting accumulation of malonyl-CoA in the mitochondria strongly inhibited carnitine palmitoyltransferase 1A (CPT1A), a key rate-limiting enzyme in fatty acid oxidation **(Fig 6g)**. In our validation experiments, we observed that in the presence of 5 μM MCD inhibitor CBM-301940, although partial cellular activity was suppressed in keratinocytes, red light irradiation still induced a significant increase in cellular activity. This indicates that the red light-driven increase in cellular activity is not mediated by enhanced lipid metabolism **(Fig 6h)**, but rather by transient effects mediated by photosensitive molecules within the cell, such as cytochrome c oxidase, in response to red light. However, under inhibitor conditions, red light no longer promoted fatty acid uptake by keratinocytes **(Fig 6i)**, and the characteristic red-light-driven increases in ATP and NADH levels in keratinocytes disappeared (**Fig 6j** and **k)**. Acetyl-CoA content and H3K9ac levels markedly decreased in keratinocytes **(Fig 6l, m**, and **n)**, and red light irradiation failed to reverse H3K9ac levels (**Fig 6m** and **n)**. These findings confirm our hypothesis that, following the regulation of keratinocyte metabolism by red light, red light maintains the acetylation of MCD by downregulating SIRT4 levels. This activates MCD function, thereby blocking the inhibitory signals on fatty acid oxidation.

Therefore, when the MCD function is inhibited, red light can no longer achieve PBM, which promotes mitochondrial fatty acid oxidation and the TCA cycle while elevating acetyl-CoA levels in keratinocytes by regulating MCD activity via SIRT4. As acetyl-CoA is the sole acetylation substrate for H3K9ac, this ultimately leads to increased H3K9ac levels in keratinocytes. This mechanism explains the anti-aging effects of red light on the skin observed in our mouse model of natural aging **(Fig 1)**. These findings conclusively demonstrate that the capacity of red light to regulate cellular lipid metabolism and restore acetyl-CoA and H3K9ac levels operates through the SIRT4-MCD axis.

### 7. Red light alleviated systemic aging in a mouse model of natural aging

Through metabolic characterization and mechanism exploration following red light treatment of cell and animal models, we elucidated the mechanism by which red light activates mitochondrial lipid metabolism and promotes H3K9ac by inhibiting the SIRT4-MCD axis. In mouse models, we validated the tissue penetration depth of red light by measuring its intensity after traversing the mouse skin using photometers, as shown in **Fig S7a** and **b**. We found that at a light source height of 10 cm, 51.04% of the red light fully penetrated the skin tissue **(Fig S7c** and **d)**. This suggests that some red light may penetrate the epidermis to reach internal organs and tissues. Consequently, we assessed the SIRT4 protein expression levels in aged animal models. We observed that periodic red light treatment significantly reversed the upregulation of SIRT4 protein in central organs such as the liver, lungs, and kidneys of aged mice **(Fig 7a-d)**. In contrast, the levels of H3k9ac increased significantly following red light exposure (**Fig 7e-l**). Fluorescence levels of p-NF-κB protein markedly decreased across all tissues and organs, indicating that periodic systemic red light irradiation not only alleviates inflammatory aging in skin tissue but also exerts anti-inflammatory effects on tissues and organs within its penetration depth range **(Fig 7m-p)**. Additionally, we assessed P16 and LAMB1 protein expression levels, along with SASP inflammatory factor expression, in heart, liver, lung, and kidney tissues. Following periodic red light treatment, P16 protein expression decreased, whereas Lamin-B1 protein expression increased **(Fig 7 m-p)**. The levels of most aging-associated SASP inflammatory factors significantly decreased after periodic red light treatment **(Fig 7q-t)**. These findings further demonstrate, in an animal model, that periodic red light irradiation possesses tissue anti-aging effects.

**Figure 7.**
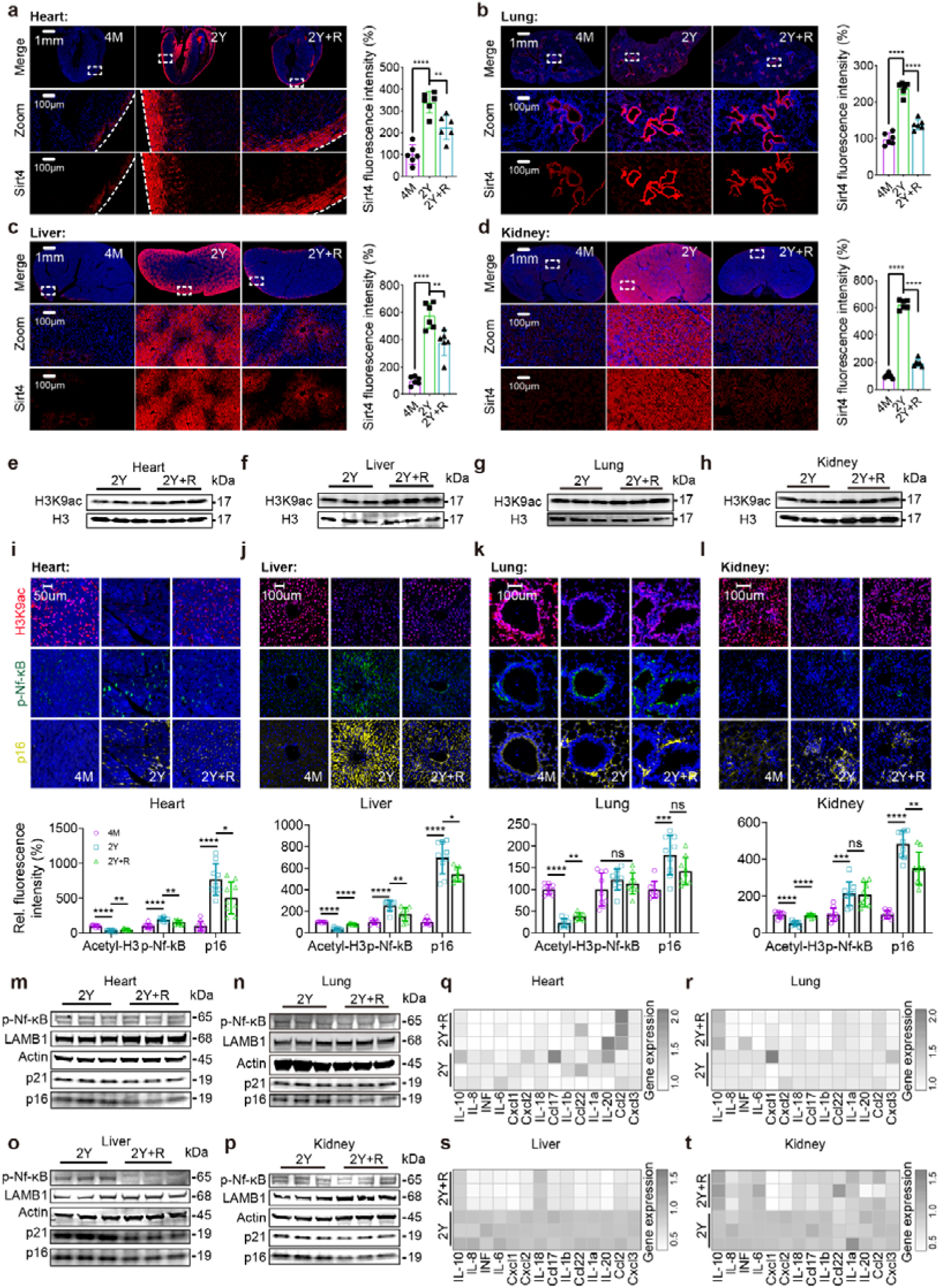
Inflammation and senescence signaling in aging model mice alleviated after cyclic treatment with red light. **a, b, c** and **d,** Fluorescence images of SIRT4 (red) proteins in the heart (**a**), liver (**b**), lung (**c**)and kidney (**d**) of mice in groups 4M, 2Y and 2Y+R (DAPI, blue), n=6. **e, f, g** and **h,** The levels of H3K9ac proteins in the heart (**e**), liver (**f**), lung (**g**)and kidney (**h**) of mice in groups 4M, 2Y and 2Y+R, n=3. **i, j, k** and **l,** Fluorescence images of H3K9ac (Red), p-Nf-κB (green) and p16 (yellow) proteins in the heart (**i**), liver (**j**), lung (**k**)and kidney (**l**) of mice in groups 4M, 2Y and 2Y+R (DAPI, blue), n=6. **m, n, o** and **p,** The levels of p-Nf-κB, LAMB1, p21 and p16 proteins in the heart (**m**), liver (**n**), lung (**o**)and kidney (**p**) of mice in groups 4M, 2Y and 2Y+R, n=3. **q, r, s** and **t,** The expression heatmap of SASP inflammatory factors in the heart (**q**), liver (**r**), lung (**s**)and kidney (**t**) of mice in groups 4M, 2Y, and 2Y+R, n=3. *P* values were calculated applying an unpaired Student’s *t*-test with unequal variances. ns = not significant (*P* > 0.05).

In summary, our findings demonstrate that red light can reverse cellular senescence at both the cellular and tissue levels by downregulating SIRT4 protein expression. By elucidating the mechanism through which red light upregulates H3K9ac protein levels, we confirmed its potential role in mitigating cellular and tissue senescence by enhancing fatty acid oxidation and mitochondrial metabolism via SIRT4 downregulation, thereby increasing H3K9ac. We propose that SIRT4 protein downregulation represents a novel anti-aging intervention mechanism that activates mitochondrial and fatty acid metabolism to elevate H3K9ac levels and ultimately mitigate cellular aging. By controlling the abnormal age-related activation of SIRT4 during cellular aging processes and developing SIRT4 inhibitors, tissue-specific aging reversal and therapeutic interventions for metabolic-related diseases may be realized.

## Discussion

The sirtuin family of proteins is also known as the longevity protein family^28^. SIRT1, SIRT3, and SIRT6 regulate multiple aging-related pathways by linking NAD^+^-dependent deacetylase activity to cellular energy metabolism, stress resistance, inflammation, and protein acetylation.^30, 49, 52–54^. In contrast, mitochondrial SIRT4 appears to exert functions that partially oppose those of other sirtuins. SIRT4 activation can inhibit fatty acid oxidation and mitochondrial metabolism and has also been associated with reduced MAPK signalling^55, 56^. Hepatic SIRT4 knockout has been reported to enhance SIRT1 expression and restore PPAR-α-mediated lipid metabolism, suggesting that SIRT4 may function as a feedback regulator within the sirtuin network^51^. Consistent with this concept, our study showed that SIRT1 progressively declined in the mouse skin stratum corneum during aging, whereas SIRT4 increased **(Fig 5h)**. Periodic red light treatment reversed this age-associated pattern **(Fig 5d-f)**, and the molecular and metabolic changes induced by red light occurred predominantly in keratinocyte mitochondria. These observations are important because keratinocytes constitute a major epidermal cell population directly exposed to therapeutic red light and can transmit metabolic and inflammatory signals to surrounding skin compartments. Thus, changes in keratinocyte SIRT4 may provide a cellular entry point through which a noninvasive optical stimulus reshapes tissue aging. Together, our findings support a model in which red light modulates skin aging by reducing SIRT4-mediated metabolic restraint in keratinocytes.

Similarly, histone acetylation is closely linked to cellular metabolism^19^. H3K9ac, a typical transcriptional activation marker within the cellular genome, maintains active gene expression in young cells^57–59^. During aging, H3K9ac may contribute to senescence through both global loss and local abnormal accumulation^60^. On the one hand, age-associated oxidative stress, reduced acetyltransferase activity, and impaired mitochondrial function decrease acetyl-CoA availability and thereby reduce overall histone acetylation^20^. On the other hand, inflammatory signals such as TNF-α can activate IKK and phosphorylate p65, leading to local H3K9ac enrichment at SASP inflammatory gene loci, including IL-6 and IL-8^61–64^. In our study, SIRT4 downregulation activated mitochondrial metabolic processes, especially fatty acid metabolism and the TCA cycle, through enhanced acetylation of mitochondrial metabolism-related proteins such as MCD. This was accompanied by increased acetyl-CoA accumulation in SIRT4-downregulated cells **(Fig 6)** and restoration of H3K9ac levels in aging skin. At the same time, reduced SIRT4 expression restored SIRT1 protein levels in aged cells and tissues^65–67^. Therefore, although the overall consequence of SIRT4 downregulation was increased H3K9ac, we also observed remodeling of SASP factor expression and attenuation of p-NF-κB-related inflammatory signalling **(Fig 7)**. This apparent dual effect is consistent with the concept that histone acetylation is not simply beneficial or harmful, but depends on genomic location, metabolic context, and inflammatory state. Nevertheless, the inflammatory effects of red light should not be interpreted as uniform suppression of all cytokines. Red-light-induced metabolic activation may reduce the senescence-associated inflammatory baseline while allowing transient reparative or stress-related cytokine responses. This interpretation may explain why some cytokines can rise in specific short-term settings while the broader aging-associated inflammatory phenotype is alleviated after periodic treatment. Thus, red light-driven SIRT4 downregulation may remodel senescent gene expression by coordinating global H3K9ac restoration, SIRT1-associated inflammatory control, and mitochondrial metabolic recovery.

Additionally, we observed that the most significant cellular change following 625-635 nm red light irradiation of keratinocytes was a red light-driven metabolic promotion. This effect was distinctly different from our previous findings showing that blue-light irradiation reduced keratinocyte viability in a dose-dependent manner^37^. This wavelength-dependent difference suggests that the biological effects of red light should be considered from the perspective of cellular metabolic reprogramming. Existing research suggests that red light may be detected by the mitochondrial photoreceptor system (including cytochrome c oxidase)^40, 68^, and current findings largely support the notion that red light regulates mitochondrial metabolism. After entering cells as an energy form, red light may help maintain mitochondrial membrane potential and alter mitochondrial efficiency, thereby changing the balance of intracellular metabolites^69, 70^. After entering cells as an energy form, red light may help maintain mitochondrial membrane potential and alter mitochondrial efficiency, thereby changing the balance of intracellular metabolites. Such metabolite changes could activate metabolite-sensing pathways and reshape protein stability^56, 71, 72^. For example, metabolic activation can promote oxidation of methionine or cysteine residues, expose E3 ligase-binding motifs, and trigger ubiquitin-dependent degradation^73^. Glycosylation may also influence protein stability by competing with phosphorylation at regulatory sites^74^. These mechanisms suggest that red light may reduce SIRT4 protein levels through metabolism-dependent protein turnover, although this possibility requires further investigation. Future studies focused on SIRT4 ubiquitination, protein half-life, and mitochondrial proteostasis will help explain how a red-light signal is translated into reduced SIRT4 abundance.

A recent study by Herrera et al. provides direct evidence that red-light irradiation stimulates mitochondrial fatty acid oxidation in keratinocytes^75^. Using metabolic flux analyses, they showed that 660 nm red light enhanced basal, ATP-linked, and maximal oxygen consumption rates and that this effect was inhibited by etomoxir, supporting FAO-dependent respiration^75^. They further reported AMPK activation, ACC phosphorylation, reduced free fatty acids, and no major increase in electron transport chain protein abundance. Importantly, their study also showed that the red-light-induced increase in oxygen consumption persisted beyond the immediate irradiation period, supporting the idea that red light initiates durable metabolic reprogramming rather than only a transient photochemical effect^75^. These findings complement our results by independently supporting red-light-induced lipid metabolic activation. Our data extend this framework by identifying SIRT4 downregulation and MCD acetylation as a potential mitochondrial axis that releases fatty acid metabolic suppression, increases acetyl-CoA availability, and promotes H3K9ac restoration. The SIRT4-MCD axis and AMPK/ACC pathway may therefore represent parallel or convergent mechanisms of red-light-induced fatty acid metabolic activation. In this integrated model, AMPK/ACC signalling may rapidly increase fatty acid entry into oxidative pathways, whereas SIRT4 downregulation may sustain mitochondrial enzyme acetylation and acetyl-CoA production. Future work should test whether red light-induced SIRT4 downregulation causally affects AMPK/ACC phosphorylation and FAO-dependent respiration.

Finally, Several limitations should be considered. First, although pharmacological experiments support the involvement of PI3K/Akt/mTOR-related metabolic signalling, these results should be interpreted as supportive rather than definitive causal evidence. Genetic perturbation of PI3K/Akt/mTOR components and physiological activation with insulin or EGF would further strengthen the pathway model. Second, isolated mitochondria, permeabilized-cell assays, and direct wavelength comparisons are needed to clarify whether red light acts through direct mitochondrial photoreception, secondary metabolic signalling, or both. These experiments would also help distinguish red-light-specific effects from responses caused by light exposure in general. Third, this study used female C57BL/6 mice only; future studies should include both sexes to determine whether sex-dependent aging and metabolic responses influence red-light sensitivity.

In summary, our study reports a red-light-mediated anti-aging mechanism in which reduced SIRT4 protein levels release mitochondrial metabolic suppression in senescent keratinocytes, restore PPAR-α-regulated lipid metabolism, increase acetyl-CoA availability, and enhance aging-associated histone acetylation. Together with the direct FAO evidence from Herrera et al., our findings place SIRT4 within a broader red-light metabolic framework connecting mitochondrial lipid metabolism, acetyl-CoA homeostasis, H3K9ac remodeling, and inflammatory restraint. These results distinguish SIRT4 from other sirtuin family members and suggest that controlling age-associated SIRT4 activation may represent a potential strategy for anti-skin-aging intervention. Identifying SIRT4 inhibitors or red-light-based approaches that safely modulate SIRT4 may provide molecular guidance for future development of anti-aging therapies and skin-repair products.

## Acknowledgement

This work was supported by the National Natural Science Foundation of China (31971315) and the Postdoctoral Science Foundation of China (2024M754229). We are also grateful to Wu Si for her substantial support of this research.

## Author contributions

Fangqing Deng: design, acquisition of data, analysis and interpretation of data conceptualization, review & editing, methodology, investigation, data curation, and writing—original draft. Yingchun Yang and Zhaoxiang Yu: resources and writing–reviewing. Xu Li, Zibo Gao, Lihua Yang, Huifang Liu, Rong Yang, Monian Wang, Yang Liu, Jinyun Niu: investigation. Lianbing Zhang: conceptualization, review & editing, funding acquisition, resources, and supervision.

## Data availability

All data are available in the main text or the Supplementary Materials. Original gel images of the western blots used in this study are shown in Supplementary Figs. 8-9. The RNA sequencing data related to this study has been deposited into the Sequence Read Archive (SRA) database, with accession code http://www.ncbi.nlm. nih.gov/bioproject/1161480. The Proteomics data and Acetylated proteomics data related to this study has been deposited into the The Genome Sequence Archive (GSA) database, with accession code https://ngdc.cncb.ac.cn/omix/select-edit/OMIX011505 and https://ngdc.cncb.ac.cn/omix/select-edit/OMIX011504.

