## Supplementary materials for "Photo-downregulation of SIRT4 mitigates aging in mice by enhancing H3K9ac via fatty acid metabolism"

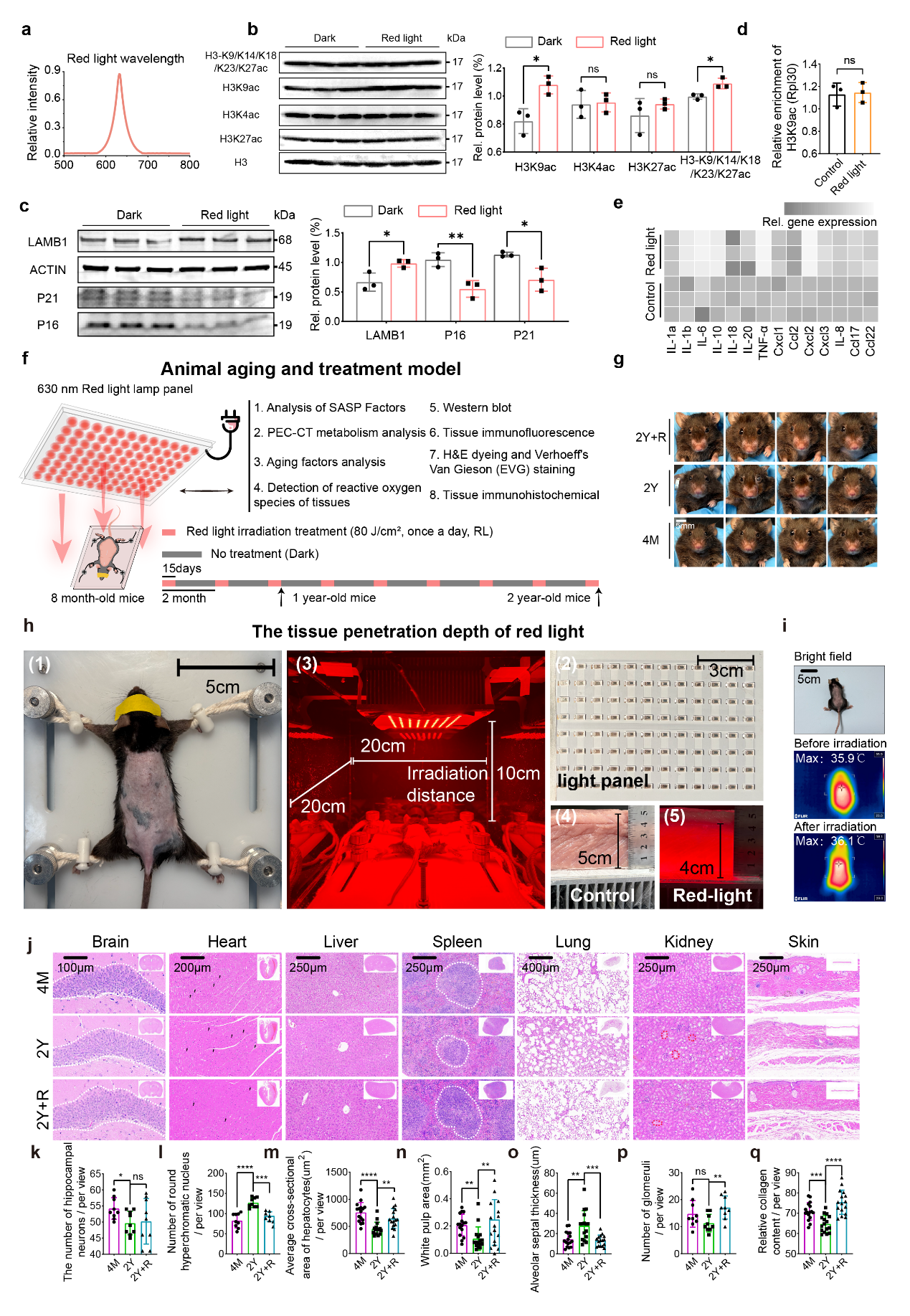

**Fig S1. a,** The wavelength range of the red LED lamp used in this study. **b,** Protein expression levels of H3K4ac, H3K27ac, H3K9ac, H3-K9/K14/K18/K23/K27ac, and H3 in keratinocytes treated with red light. n=3. **c,** Protein expression levels of P16, P21, Actin and LAMB1 in aging keratinocytes treated with red light, n=3. **c,** Expression heatmap of SASP inflammatory factors in aging keratinocytes. n=3. **d,** The effect of red light irradiation on the level of H3K9ac enrichment at the RPL30 locus, n=3. **e,** Schematic diagram of animal senescence and red light irradiation treatment protocols. Description of the irradiation device used for aging C57 mice. Female C57 mice up to 8 months of age were used for periodic red light irradiation. Every two months, the mice were irradiated for 15 days at a daily dose of 80 J/cm^2^. As shown in a, one-year-old mice were irradiated three times for 15 days, whereas two-year-old mice were irradiated nine times for 15 days. After the red light irradiation program, one- and two-year-old mice were used for subsequent analysis. **f**, Facial images of mice at different ages after cyclic red light irradiation. n=6. **g**, Light protocol and tissue penetration depth validation of the red light LED device used in this study. As shown in **(1)**, eye-covered mice were immobilized on a stand and irradiated using the red lamp panel shown in **(2)** and **(3)**. Visible light penetration experiments were performed using muscle tissue with a thickness of 5 cm to simulate mouse skin tissue, as shown in **(4)** and **(5)**. The results showed that the tissue penetration of the red light could reach up to 4 cm under this condition. **h,** Infrared imaging of mice following red light irradiation. **i,** H&E staining of the brain, heart, liver, lungs, kidneys, spleen and skin of mice in the 4M, 2Y and 2Y+R treatment groups. **j-p,** Number of hippocampal nerve cells (**j**, n=9), number of hyperchromatic nuclei (**k**, n=9), average cross-sectional hepatocytes (l, n=18), white pulp area (**m**, n=18), alveolar septal thickness (**n**, n=18), number of glomeruli (**o**, n=9), and relative collagen content of the skin (**p**, n=18) in the H&E staining field of view of tissues from the 4M, 2Y, and 2Y+R treatment groups.

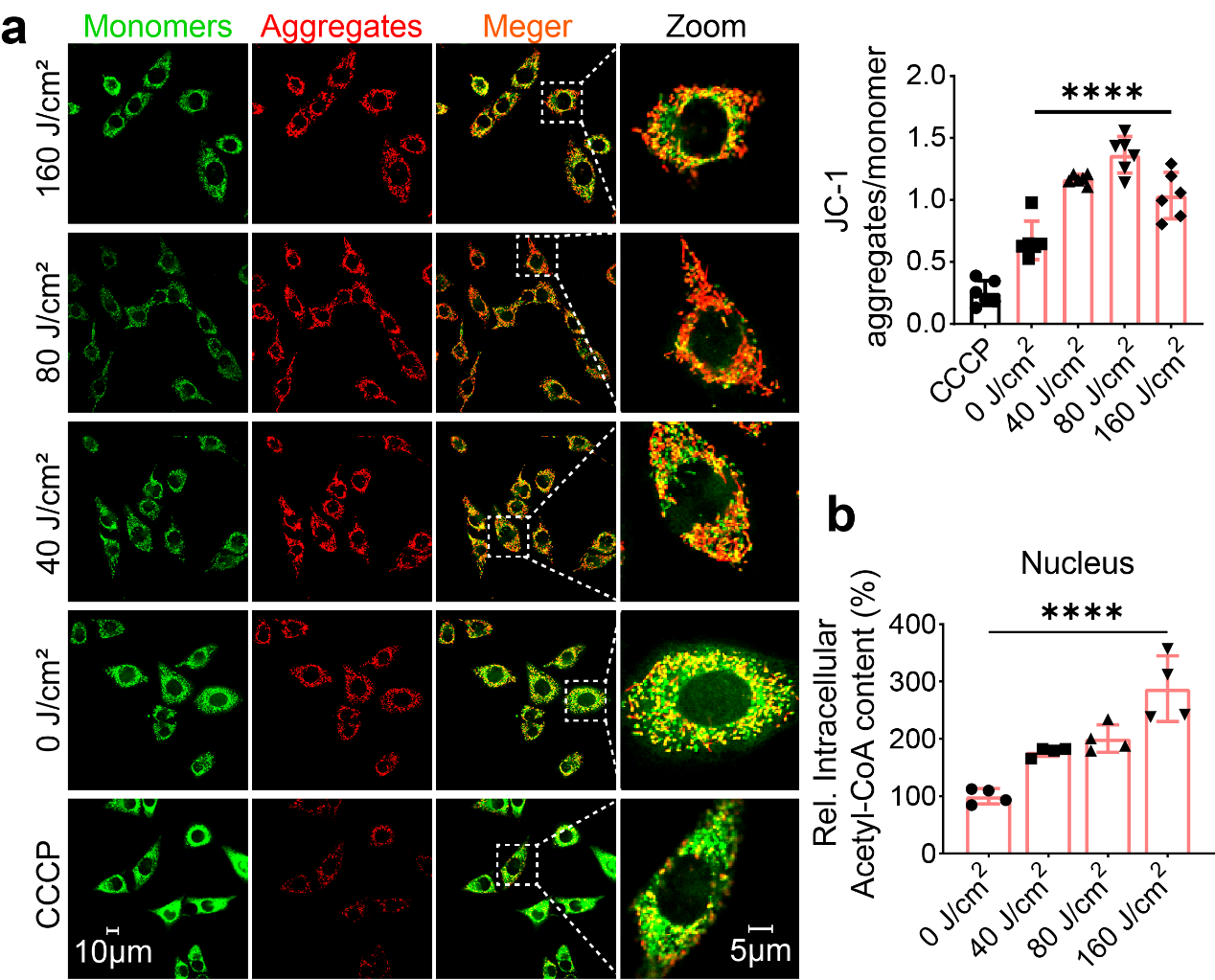

**Fig S2. a,** Mitochondrial membrane potential images of keratinocytes under different red light doses. **b,** Relative changes in nucleus acetyl-CoA of keratinocytes with graded doses of red light, n=4.

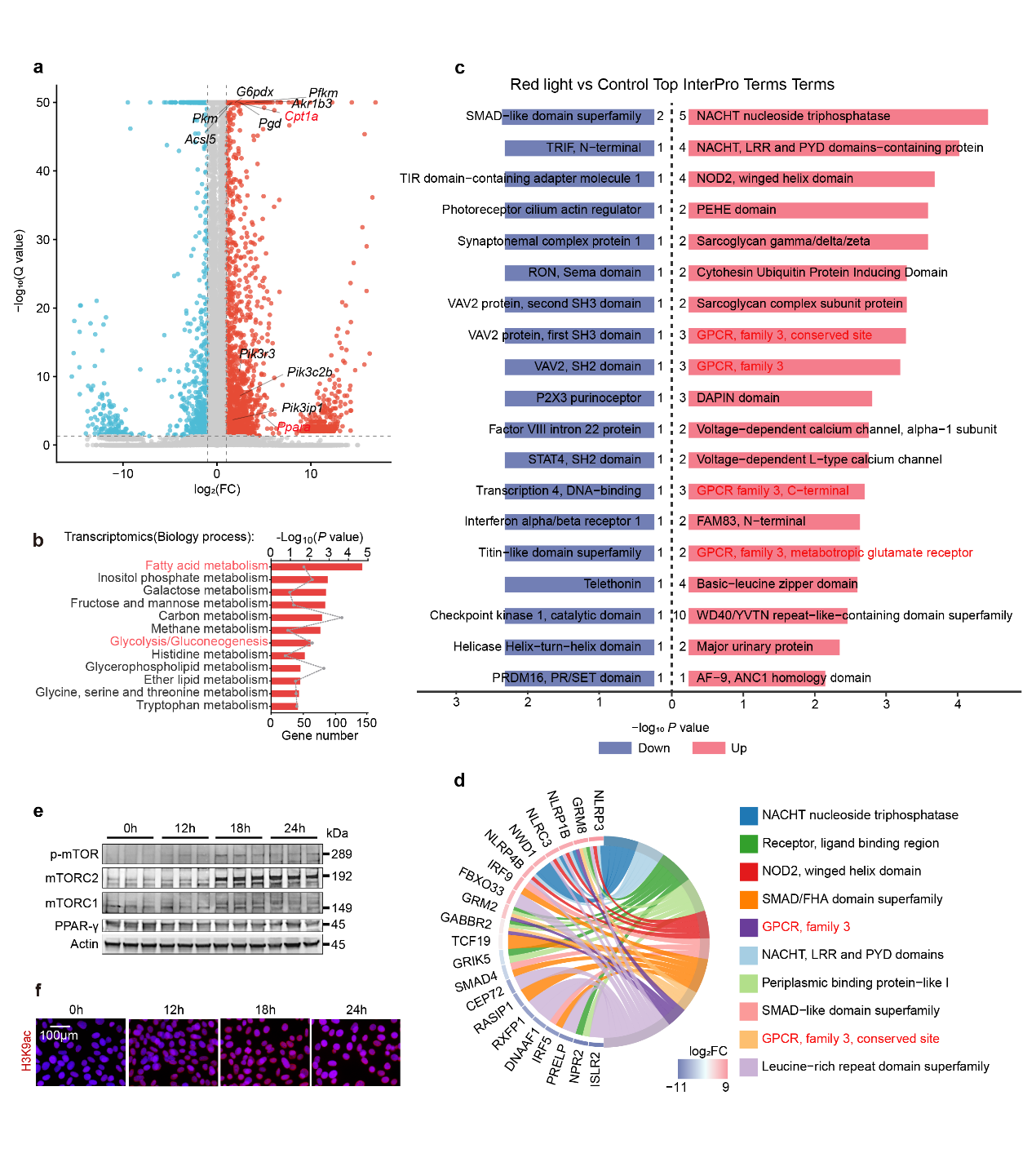

**Fig S3. a,** Volcano plot of the differences in gene expression after red light treatment. Conducting a significance test using the corrected p-value (Q-value). Volcano plot showing differentially expressed genes between the red-light irradiation and control groups. Differentially expressed genes were defined as Q value < 0.05 and |log2 fold change| > 1. For visualization, Q values smaller than 1 × 10^-50^ were capped at 1 × 10^-50^. Red and blue dots indicate significantly upregulated and downregulated genes, respectively; grey dots indicate non-significant genes. **b,** Enrichment analysis of metabolic pathways after red light treatment (P value > 0.5). **c** and **d,** Proteomic analysis of red light-irradiated keratinocytes, showing the pathways enriched for significant differences in Top 20 up- and down-regulation by Inter Pro analysis (**c**) and chord diagrams of Top 10 processes with significant differences obtained from Inter Pro analysis (**d**). **e,** Changes in the levels of p-mTOR, mTORC1, mTORC2, PPAR-γ proteins in keratinocytes at 0h, 12h, 18h, and 24h after red light treatment, n=3. **f,** Fluorescence images of H3K9ac protein at different time points after red light treatment in keratinocytes.

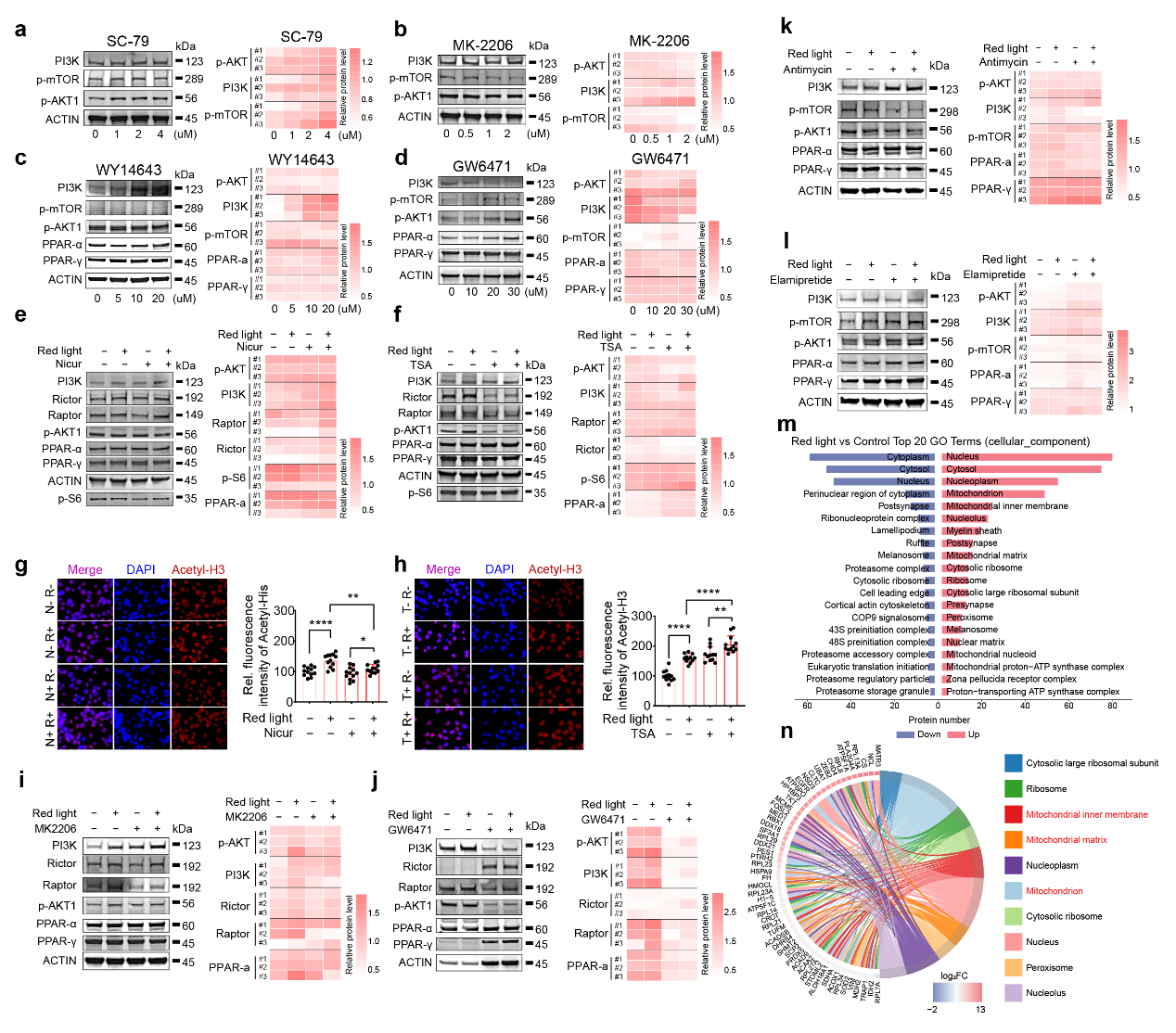

**Fig S4. a** and **b,** Effects of different concentrations of SC-79 (**a**) and MK2206 (**b**) levels of PI3K, p-mTOR, p-AKT and Actin proteins in keratinocytes, n=3 (DAPI, blue). **c** and **d,** Effects of different concentrations of WY14643 (**c**) and GW6471 (**d**) levels of PI3K, p-mTOR, p-AKT, PPAR-α, PPAR-γ and Actin proteins in keratinocytes, n=3 (DAPI, blue). **e** and **f,** Effect of red light treatment on the levels of PI3K, Raptor, Rictor, p-AKT, PPAR-α, PPAR-γ and p-S6 proteins in keratinocytes in the presence of 5uM Nicur (**e**) and TSA (**f**), n=3. **g** and **h,** Fluorescence images of intracellular H3K9ac protein in keratinocytes treated with red light irradiation in the presence of 5uM Nicur (**g**) and 5uM TSA (**h**). **i** and **j,** Effect of red light treatment on the levels of Pi3k, Raptor, Rictor, PPAR-α, PPAR-γ, p-S6 and Actin proteins in keratinocytes in the presence of 2uM Mk2206 (**i**) and 20uM GW6471 (**j**), n=3. **k** and **l,** Effect of red light treatment on the levels of PI3K, p-mTOR, p-AKT, PPAR-α, PPAR-γ and Actin proteins in keratinocytes in the presence of 5uM Antimycin (**k**) and Elamipretide (**l**), n=3. **m** and **n,** Acetylation modification proteomics analysis illustrating differential expression GO analysis (cellular component pathway, **m**) and chordal analysis (**n**) of acetylation-modified proteins in cells after red light irradiation of keratinocytes.

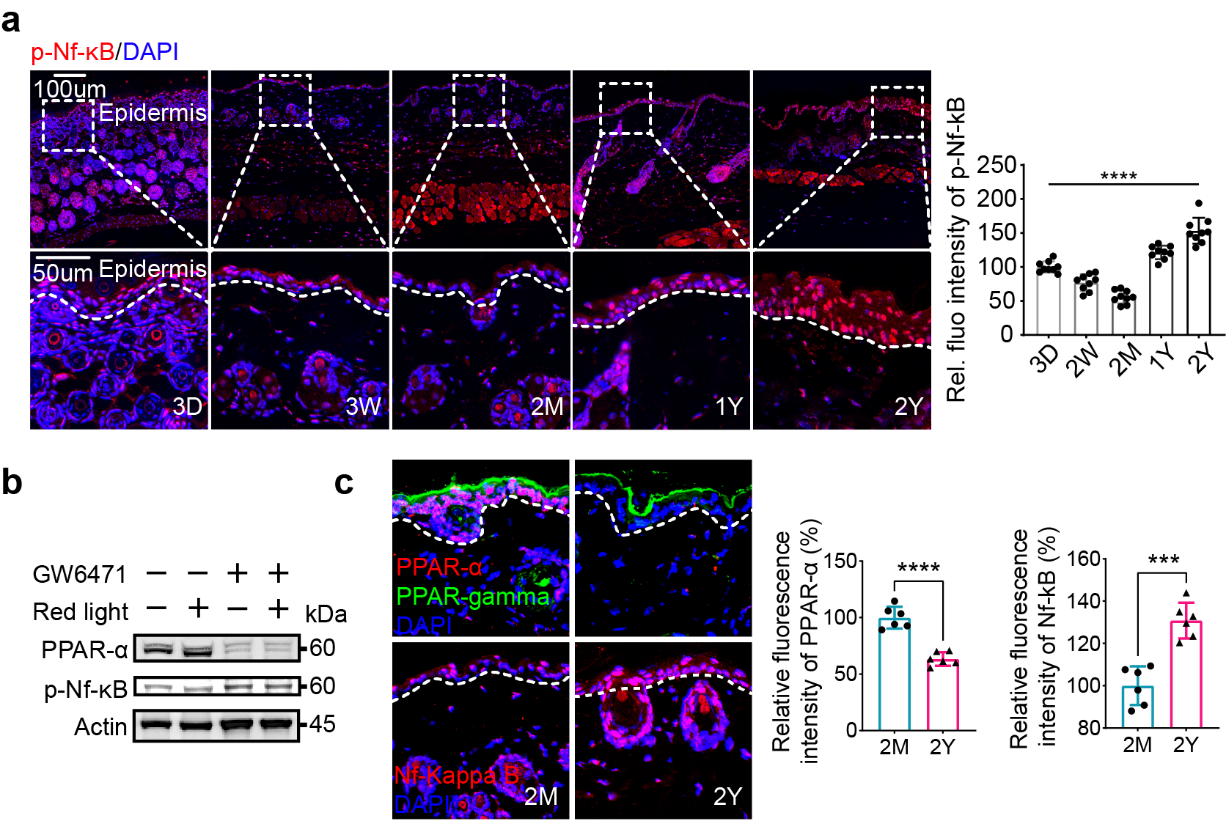

**Fig S5. a,** Fluorescence images of p-NF-κB protein (p-NF-κB, red; DAPI, blue) in skin tissues from 3D, 3W, 2M, 1Y, and 2Y mice. **b,** Intracellular protein levels of PPAR-α and p-NF-κB after RL treatment in the presence of the PPARα inhibitor GW6471 (20 µM). **c,** Fluorescent images of PPAR-α, PPAR-γ, and p-NF-κB in the skin of 2M and 2Y old mice (DAPI, blue).

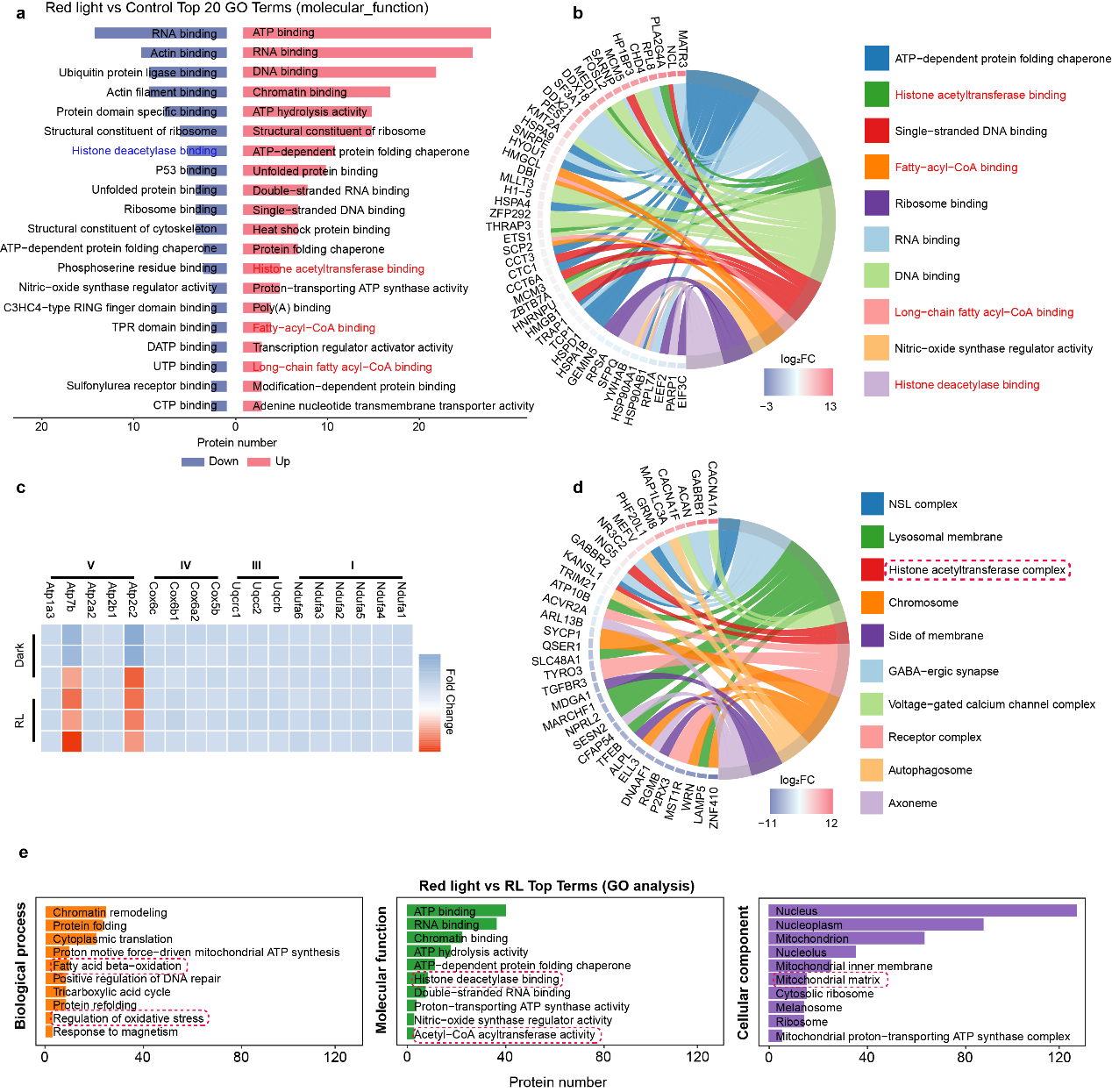

**Fig S6. a** and **b,** Acetylation modification proteomics analysis, illustrating differential expression GO analysis (molecular function pathway, **a**) and chordal analysis (**b**) of acetylation modification proteins in cells after red light irradiation of keratinocytes. **c,** Heat map of changes in the content of mitochondrial electron-conducting chain complex-related proteins in keratinocytes after red light treatment. **d,** Proteomic and chordal plots after red light treatment of keratinocytes, demonstrating the processes involved in the significant difference Top10 in the Cellular component pathway. **e,** Conjoint GO analysis of proteomics and acetylation modification proteomics after red light treatment of keratinocytes demonstrated significant differences Top10 processes in Biological process, Cellular component and Molecular function pathways within keratinocytes, respectively.

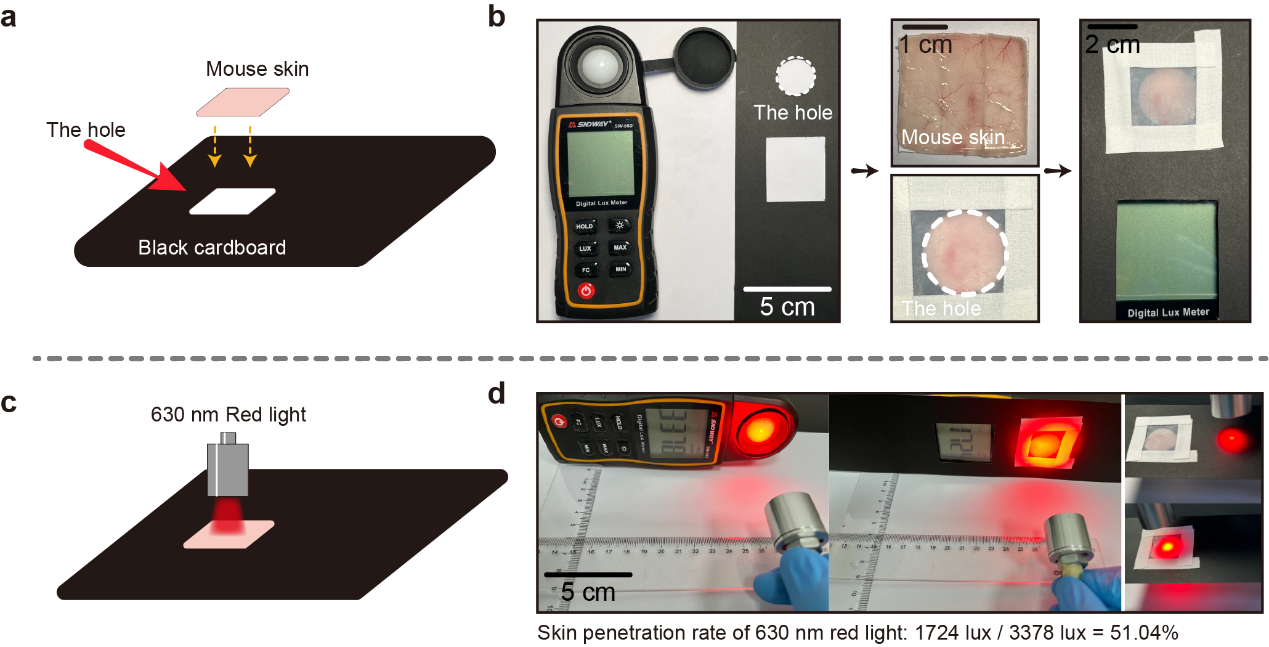

**Fig S7. a** and **b,** Schematic diagram for testing the penetration of red light through the skin **(a)**. An illuminometer was used to detect the light intensity of the LED source before and after penetrating the skin **(b)**. **c** and **d,** Schematic diagram illustrating the measurement of red light penetration efficiency in skin tissue **(c)**.The skin penetration efficiency of the LED source is represented by the ratio of the illuminometer readings under direct illumination and illumination through the skin at the same distance for visible red LED sources **(d)**.

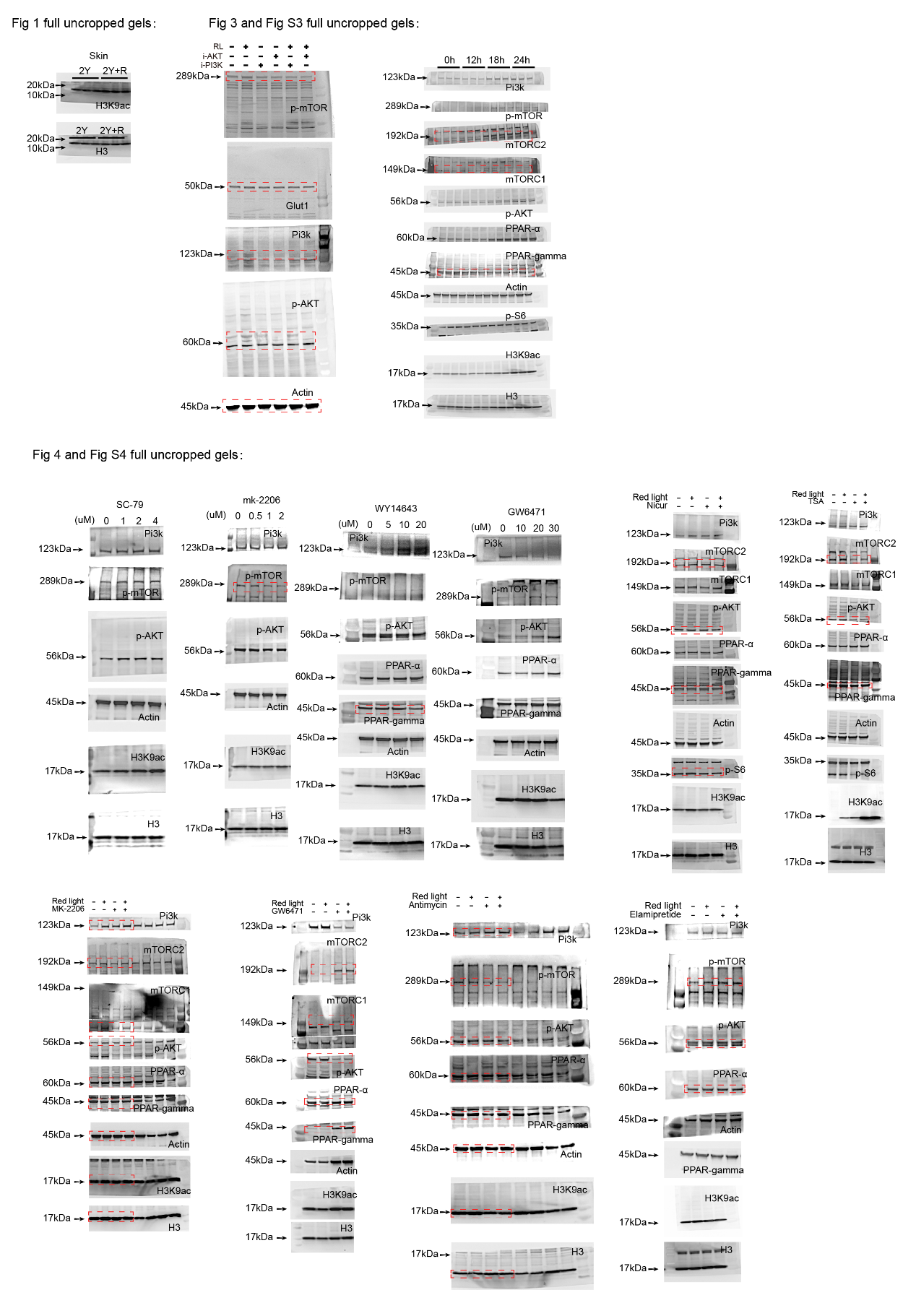

**Fig S8.** Uncropped gels for Western Blots in Figure 1, Figure 3 and Figure 4.

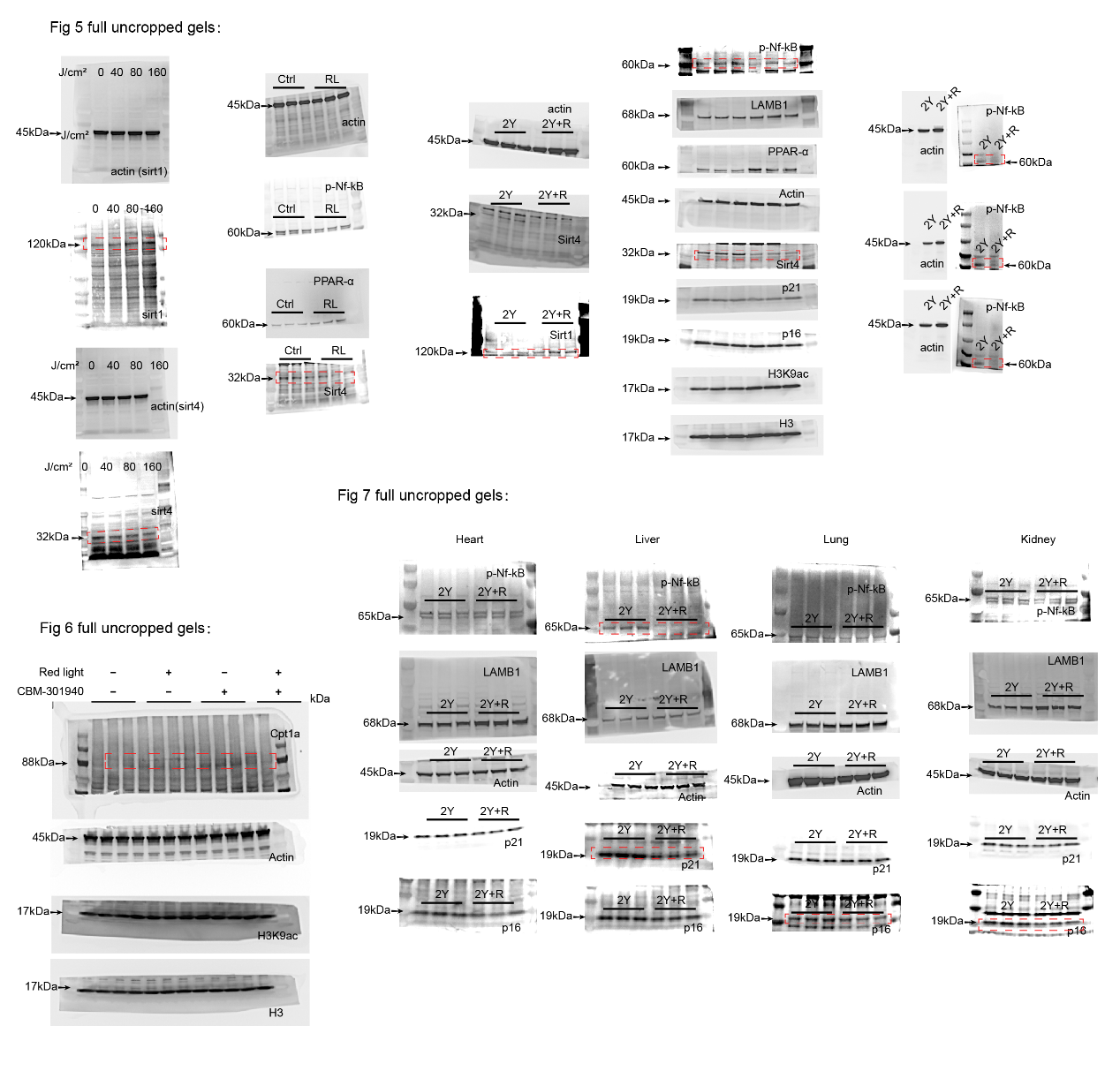

**Fig S9.** Uncropped gels for Western Blots in Figure 5, Figure 6 and Figure 7.

Table S1: qPCR primers used in this work.

| Table S1:qPCR primers used in this work. | | |
| --- | --- | --- |
| **Gene Names** | **Forward primer** | **Reverse primer** |
| ***IL-10*** | ATGCTGCCTGCTCTTACTGACTG | CCCAAGTAACCCTTA AAGTCCTGC |
| ***Cxcl3*** | CCAGACAGAAGTCATAGCCAC | CTTCATCATGGTGAGGGGCTT |
| ***IL-1α*** | CGCTTGAGTCGGCAAAGAAAT | TGGCAGAACTGTAGTCTTCGT |
| ***IL-1b*** | GCCACCTTTTGACAGTGATGAG | GACAGCCCAGGTCAAAGGTT |
| ***IL-6*** | GACAAAGCCAGAGTCCTTCAGA | GTGACTCCAGCTTATCTCTTGG |
| ***IL-18*** | ACAAGCATCCAGGCACAGC | AAGGTTTGAGGCGGCTTTCT |
| ***Ccl2*** | AGATGCAGTTAACGCCCCAC | GAGCTTGGTGACAAAAACTACAGC |
| ***IL-20*** | TCGGGATAGTGTGTCTTTGGA | GCGAGGCTGCTGATCTTTCT |
| ***TNF-α*** | CCCTCACACTCACAAACCAC | ACAAGGTACAACCCATCGGC |
| ***Cxcl1*** | CCCAAACCGAAGTCATAGCCA | CCGTTACTTGGGGACACCTT |
| ***Cxcl2*** | CCAAAAGATACTGAACAAAGGCAAG | ATCAGGTACGATCCAGGCTTC |
| ***IL-8*** | GGCCCAATTACTAACAGGTTCC | TCTCTTGTTCTCAGGTCTCCCA |
| ***Ccl22*** | GAGATCTGTGCCGATCCCAG | CCACGGTCATCAGAGTAGGC |
| ***Ccl17*** | AATGTAGGCCGAGAGTGCTG | GACAGTCAGAAACACGATGGC |

Table S2: ChIP-qPCR primers used in this work.

| Table S2:ChIP-qPCR primers used in this work. | | | |
| --- | --- | --- | --- |
| Gene Names | | Forward primer | Reverse primer |
| **SOD2** | -208/-353 | ggtgaaggtgcagccatagt | aagggctggcatcacttctc |
|  | -1745/-1861 | aggggccctgattactccat | gtgagctgcaaagcttccac |
|  | -1078/-1238 | ctgccatgctcccaccttaa | tcagaccggagctgtatgga |
| **Cdkn2a(P21)** | -1908/-2107 | gtctgttcagtcctgggtgg | ggcaaagtgggacgtcctta |
|  | -256/-449 | ggctcacttacagttccccc | gacactctgctccacacaca |
|  | -1684/-1892 | tcccctgtccttttctggga | tccgattttgctgctggtct |
| **Cdkn2a(P16)** | -1933/-2072 | caggtcaggagcagagtgtg | gatggctctcctcgagttcg |
|  | -1290/-1480 | ttgccctccgatgacttcac | agctacaacttcgccaagct |
|  | -1022/-1238 | aggcagaagggagacagagt | ttggtgtgctgcgcataaac |
| **PGC-1α** | -1502/-1783 | ggggtgttgccttcaaacac | gccaccaactctaaaccgga |
|  | -1764/-1961 | tccggtttagagttggtggc | cctcccttctcctgtgcaag |
|  | -719/-1001 | ggtcccctgtgcatttctca | ctccaaccctagtgccttgg |
| **LAMB1** | -1346/-1501 | aacggaaaacagagggtgca | cccacttttggctgtgtgtg |
|  | -1003/-1144 | ggaaccactccacgtgtagg | ggtttttgcttcgtcacccc |
|  | -1840/-2074 | gcccgagggtcagattttga | ccgcagcactttgtttcctc |
